# PFA ependymoma-associated protein EZHIP inhibits PRC2 activity through a H3 K27M-like mechanism

**DOI:** 10.1101/585729

**Authors:** Siddhant U. Jain, Truman J. Do, Peder J. Lund, Andrew Q. Rashoff, Marcin Cieslik, Katharine L. Diehl, Andrea Bajic, Nikoleta Juretic, Shriya Deshmukh, Sriram Venneti, Tom W Muir, Benjamin A. Garcia, Nada Jabado, Peter W. Lewis

## Abstract

Polycomb group (PcG) proteins are essential for development and are frequently misregulated in human cancers. Polycomb Repressive Complexes (PRC1, PRC2) function in a collaborative epigenetic cross-talk with H3K27me3 to initiate and maintain transcriptional silencing. Diffuse intrinsic pontine gliomas (DIPGs) have extremely low H3K27me3 levels mediated by H3 K27M oncohistone. Posterior fossa type A (PFA) ependymomas also exhibit very low H3K27 methylation but lack the K27M oncohistone. Instead, PFA tumors express high levels of *EZHIP* (Enhancer of Zeste Homologs Inhibitory Protein, also termed *CXORF67*). We find that a highly conserved sequence within the C-terminus of EZHIP is necessary and sufficient to inhibit the catalytic activity of PRC2 *in vitro* and *in vivo*. Our biochemical experiments indicate that EZHIP directly interacts with the active site of the EZH2 subunit in a mechanism that is remarkably similar to the K27M oncohistone. Furthermore, expression of H3 K27M or *EZHIP* in cells promote similar chromatin profiles: loss of broad H3K27me3 domains, but retention of H3K27me3 at the sites of PRC2 recruitment. Importantly, we find that H3K27me3-mediated allosteric activation of PRC2 substantially increases the inhibition potential of EZHIP and H3 K27M, providing a potential mechanism for loss of H3K27me3 spreading from CpG islands *in vivo*. Our data indicate that PFA ependymoma and DIPG are driven in part by the action of peptidyl PRC2 inhibitors– the K27M oncohistone and the EZHIP ‘oncohistone-mimic’– that dysregulate gene silencing to promote tumorigenesis.

---

Covalent modifications to both DNA and histone proteins allow chromatin to act as a dynamic information hub that integrates diverse biochemical stimuli to regulate genomic DNA access to the transcription machinery and ultimately establish and maintain cellular phenotypes. There is increasing appreciation that aberrant chromatin modifications are involved in the pathogenesis of cancer. Nowhere is this better supported than with the groundbreaking discoveries of high-frequency, somatic mutations in histones that are drivers of oncogenesis.

The discovery of high-frequency histone H3 missense mutations (‘oncohistones’) in pediatric gliomas was the first time that histone mutations were linked to any disease ^1-3^. Since the original discovery, monoallelic missense mutations in genes encoding histone H3 have been found in a variety of solid tumors and leukemias ^4-7^. Previous studies have led to mechanistic insights into how one class of oncohistones promotes tumorigenesis by finding that lysine-to-methionine (K-to-M) substitutions transform histones from serving as substrates into specific and potent inhibitors of lysine methyltransferases. Approximately 84% of diffuse intrinsic pontine glioma (DIPG) and 60% of high-grade non-brainstem pediatric midline gliomas contain a lysine 27-methionine (K27M) mutation ^1-3^. The H3 K27M oncohistone binds to and inhibits the catalytic subunit (EZH2) of the Polycomb Repressive Complex 2 (PRC2), a conserved protein complex involved in gene silencing ^8-14^. Genome-wide profiling of H3K27me3 in H3 K27M-containing DIPG tumors and transgenic cell lines revealed a non-uniform reduction of H3K27me3 and, surprisingly, locus-specific K27M-dependent retention of this histone modification ^8,15-19^. While mechanistic questions related to H3 K27M remain, the unprecedented finding that oncohistones act as enzyme inhibitors and alter global levels of histone modifications in cells indicates a direct effect of chromatin misregulation driven by histone mutations in tumorigenesis.

A molecular subtype of ependymoma tumors exhibit extremely low H3K27 methylation levels similar to K27M-containing DIPG and midline gliomas. Ependymomas account for 10% of all pediatric tumors found in the central nervous system and can occur anywhere in the posterior fossa, spinal cord, or supratentorium ^20^. Previous studies found that posterior fossa ependymomas comprise two distinct large molecular subclasses: PFA and PFB ^21-23^. Compared to the PFB subgroup, PFA ependymomas exhibit H3K27me3 reduction and CpG island-hypermethylation similar to DIPG tumors containing the K27M oncohistone. Integrative analyses of genomic data sets from PFA and DIPG revealed that these two different tumor types share a common dysregulated chromatin landscape: global reduction of H3K27me3, but focal retention of H3K27me3 at CpG islands ^21^. Unlike the vast majority of DIPGs, only 4.2% of PFA ependymomas contain the H3 K27M mutation ^21,24,25^, and until recently it was unclear how the H3 wildtype PFA tumors achieved the aberrant DIPG-like chromatin profile.

A recent study uncovered that *CXORF67*, an uncharacterized gene whose expression is normally restricted to spermatogonia and encodes for an intrinsically disordered protein, is found at elevated levels in PFA ependymoma with poor prognosis ^26^. *CXORF67* expression was not detected in the small number of PFA ependymomas that contain the H3 K27M mutation, suggesting that these two tumor features are mutually exclusive. Additionally, CXORF67 protein co-immunoprecipitated with PRC2 subunits, and expression of *CXORF67* led to a marked reduction in H3K27me3 in cultured cells ^26^.

Here, we describe the molecular mechanism by which CXORF67 reduces H3K27me3 levels in cells. We find that CXORF67 contains a highly conserved ‘K27M-like’ sequence that is necessary and sufficient to inhibit PRC2 activity and reduce cellular H3K27me3 levels. Using isogenic cell lines, we find remarkably similar genome-wide chromatin and gene expression changes caused by expression of H3 K27M or CXORF67. Our biochemical and cell-based studies demonstrate that CXORF67 functions as a K27M-like peptidyl inhibitor of PRC2. Therefore, we propose the name EZHIP (Enhancer of Zeste Homologs Inhibitory Protein) as a more descriptive name for the function of CXORF67. Additionally, we find that EZHIP-expressing PFAs and K27M-containing DIPGs aberrantly silence the *CDKN2A* tumor suppressor gene. We conclude that two biologically and clinically related brain tumors also share a common biochemical mechanism in tumorigenesis: inhibition of PRC2 activity through expression of potent peptide inhibitors.

## RESULTS

### EZHIP forms a stable complex with PRC2 and lowers H3K27me3 *in vivo*

We sought to determine if *EZHIP* expression in ependymomas correlated with the previously noted DIPG-like chromatin profile ^21^. Using previously published RNA and ChIP sequencing datasets ^21^, we found that ependymoma tumors that express high levels of *EZHIP* also exhibit genome-wide reduction in H3K27me3 levels yet retain H3K27me3 at a subset of CpG islands (Figure S1A, B). This unique genome-wide H3K27me3 profile is remarkably similar to that observed in human DIPG tumors with the H3 K27M mutation (Figure S1A), suggesting that EZHIP and H3 K27M generate similar chromatin profiles in cells. To directly address whether EZHIP is sufficient to reduce H3K27 methylation levels, we generated human embryonic kidney-293T (HEK293T) cell lines that express FLAG epitope-tagged EZHIP, wildtype histone H3.3, H3.3 K27M or H3.3 K27R mutants. We found that expression of human EZHIP in HEK293T cell lines led to a similar overall decrease in H3K27me2/3 levels as measured by immunoblot and mass spectrometry (Figure 1A, S1D). EZHIP may promote loss of H3K27me2/3 levels in cells through direct contact and inhibition of the EZH2 subunit of PRC2, as previously demonstrated for H3 K27M oncohistone ^8,10-14^. Conversely, EZHIP may reduce H3K27me2/3 levels through disruption of the integrity of the PRC2 complex. Previous studies have found that EZH2 levels depend on the integrity of PRC2 as the loss of SUZ12 or EED subunits dramatically reduces levels of EZH2 ^27^. Importantly, we found that the steady-state levels of PRC2 subunits were not altered in cells expressing EZHIP, suggesting that the PRC2 remains intact (Figure S1C). Therefore, we suspected that EZHIP promotes loss of H3K27me2/3 by contacting and modulating the activity of PRC2.

**Figure 1:**
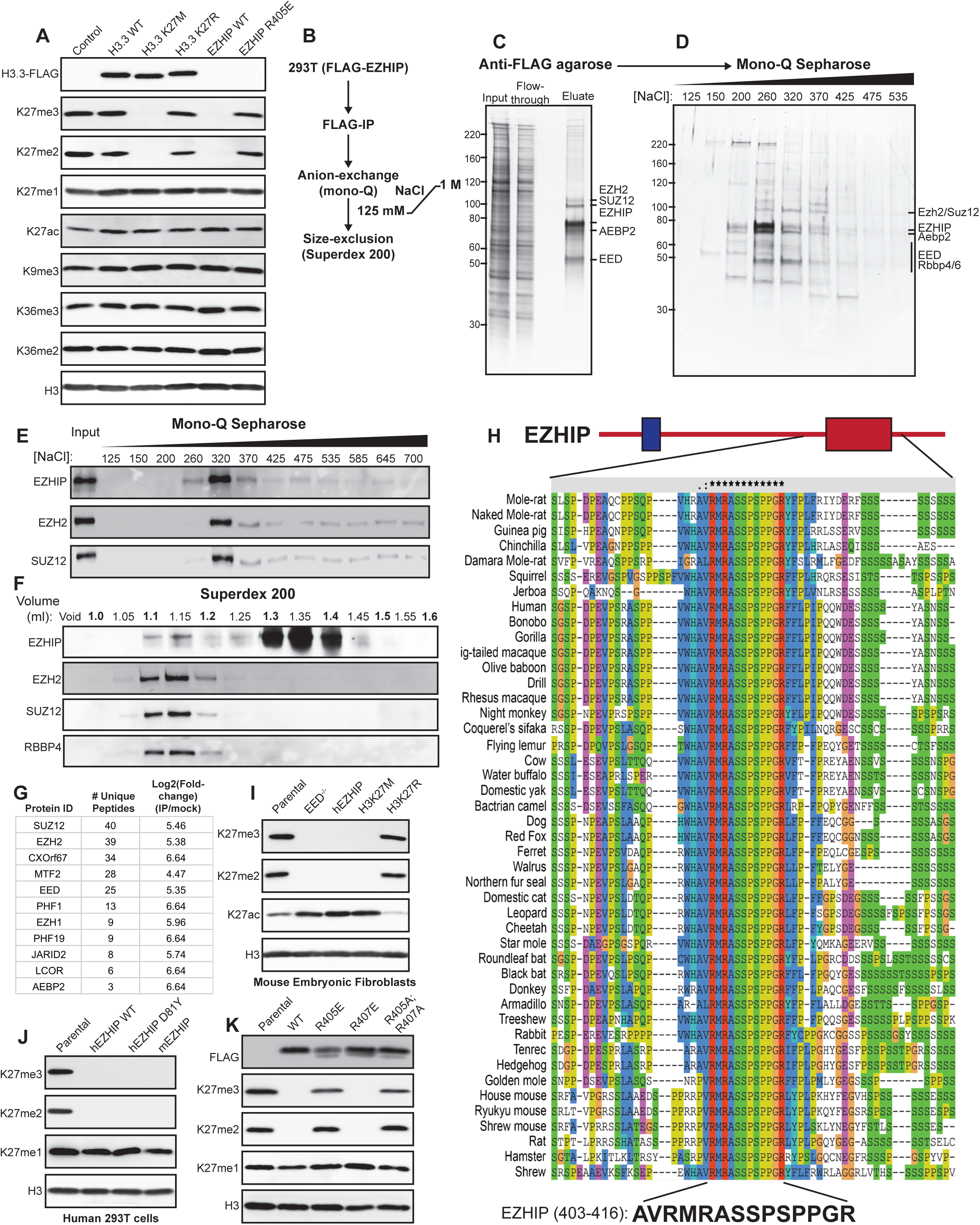
EZHIP forms a stable complex with PRC2 and lowers H3K27me3 *in vivo*. (A) Immunoblots of whole cell lysates generated from 293T cells overexpressing HA-FLAG-tagged H3.3 WT or K27M/R or FLAG-tagged EZHIP WT or R405E mutant. (B) Schematic showing the strategy for purification of EZHIP-associated proteins from 293T cells. (C-D) Silver stain of FLAG-tagged EZHIP-PRC2 complex. FLAG-EZHIP was purified from 293T cells using FLAG-immunoprecipitation (C), followed by anion-exchange chromatography (D). (E-F) Immunoblots displaying co-fractionation of EZHIP and PRC2 subunits on mono-Q and Superdex 200 columns. (G) Immunoprecipitated material from C were subjected to mass-spectrometry for protein identification and quantification. Proteins identified with at least 2 unique peptides and a log2 fold-change over the mock negative control of greater than 4 were considered hits. Protein abundances were calculated using ProteomeDiscoverer software. (H) Multiple sequence alignment displaying the conservation of EZHIP 12 amino acid sequence in the C-terminus among placental mammals. (I) Immunoblots of lysates generated from mouse embryonic fibroblasts with EED knockout or overexpressing human EZHIP WT, H3K27M or H3K27R. (J) Immunoblots of lysates generated from human 293T cell lines overexpressing human EZHIP WT or D81Y or mouse EZHIP. (K) Immunoblots of 293T overexpressing EZHIP WT, R405E, R407E or R405A;R407A mutants

To identify the EZHIP-interacting proteins in cells, we immunoprecipitated EZHIP from HEK293T cell nuclear extract using anti-FLAG M2 beads. We assigned identities to the protein bands visualized by silver stain for the M2 eluate using immunoblotting (Figure S1E and F). All core subunits of PRC2 such as EZH2, SUZ12, AEBP2, EED, and RBBP4/6 were found to associate with EZHIP (Figure 1C, S1C-E). We confirmed co-immunoprecipitation of PRC2 subunits with EZHIP using mass spectrometry for protein identification (Figure 1G). Recent studies have revealed that the core PRC2 subunits associate with mutually exclusive combinations of auxiliary subunits ^28-31^. The PRC2.1 complex includes EPOP and the PCL proteins, while PRC2.2 associates with JARID2 and AEBP2 ^30,31^. We found that EZHIP associates with both PRC2.1 and PRC2.2 as evidenced by co-immunoprecipitation of EZHIP with AEBP2, JARID2, LCOR, MTF2, PHF1, and PHF19 subunits (Figure S1E).

Immunoprecipitated EZHIP co-fractionated with PRC2 subunits on Mono Q and Superdex 200 columns (Figure 1B-F). Additionally, we found that PRC2 subunits co-immunoprecipitated with EZHIP after washing with 1.0 M KCl buffers, further demonstrating a strong association between EZHIP and PRC2 (Figure S1G). Reciprocally, FLAG-epitope tagged EZH2 immunoprecipitated endogenous EZHIP from U2OS cells (Figure S1H).

Sequence algorithms predict no stable secondary structures in EZHIP, suggesting that the protein may be intrinsically disordered. *EZHIP* homologs are exclusively found in placental mammals and, with the exception of an invariant 12 amino acid sequence near the C-terminus, show little overall sequence conservation through most of the protein (Figure 1H). Despite little overall sequence similarity, we found that expression of murine *EZHIP* in HEK293T cells, and reciprocally, human *EZHIP* in murine embryonic fibroblasts (MEFs) led to decrease in H3K27me2/3 (Figure 1I-J). These findings suggest that the conserved C-terminal sequence in EZHIP likely plays a central role in modulating PRC2 activity. Consistent with this hypothesis, a single amino acid substitution (R405E) in the conserved EZHIP peptide abolished the ability of *EZHIP* transgenes to lower H3K27me2/3 levels in HEK293T cells (Figure 1K). *EZHIP* missense mutations are found in 9.2% of PFA ependymomas and occur exclusively within a hotspot region in the poorly conserved N-terminus of protein. We found that *EZHIP* transgenes containing one of these mutations (D81Y) had no effect on H3K27me2/3 levels (Figure 1J). Similar results were recently reported for other *EZHIP* missense mutations found in PFA tumors^26^.

### EZHIP is a competitive inhibitor of PRC2

Taken together, our results suggest that EZHIP directly interacts with PRC2 and inhibits its lysine methyltransferase activity. To directly assess this proposition, we determined whether the recombinant EZHIP protein could inhibit PRC2 activity using *in vitro* methyltransferase assays. Titration of full-length, recombinant EZHIP to a methyltransferase reaction led to a dose-dependent loss of PRC2 methylation of recombinant nucleosomes (IC_50_ = 50 nM) (Figure 2A-B, S2A). Recombinant EZHIP containing the R405E substitution had a negligible effect on PRC2 activity *in vitro* (IC_50_ = NA). This finding directly links the loss of H3K27me2/3 levels in HEK293T cells with the ability of EZHIP to inhibit PRC2 activity *in vitro*.

**Figure 2:**
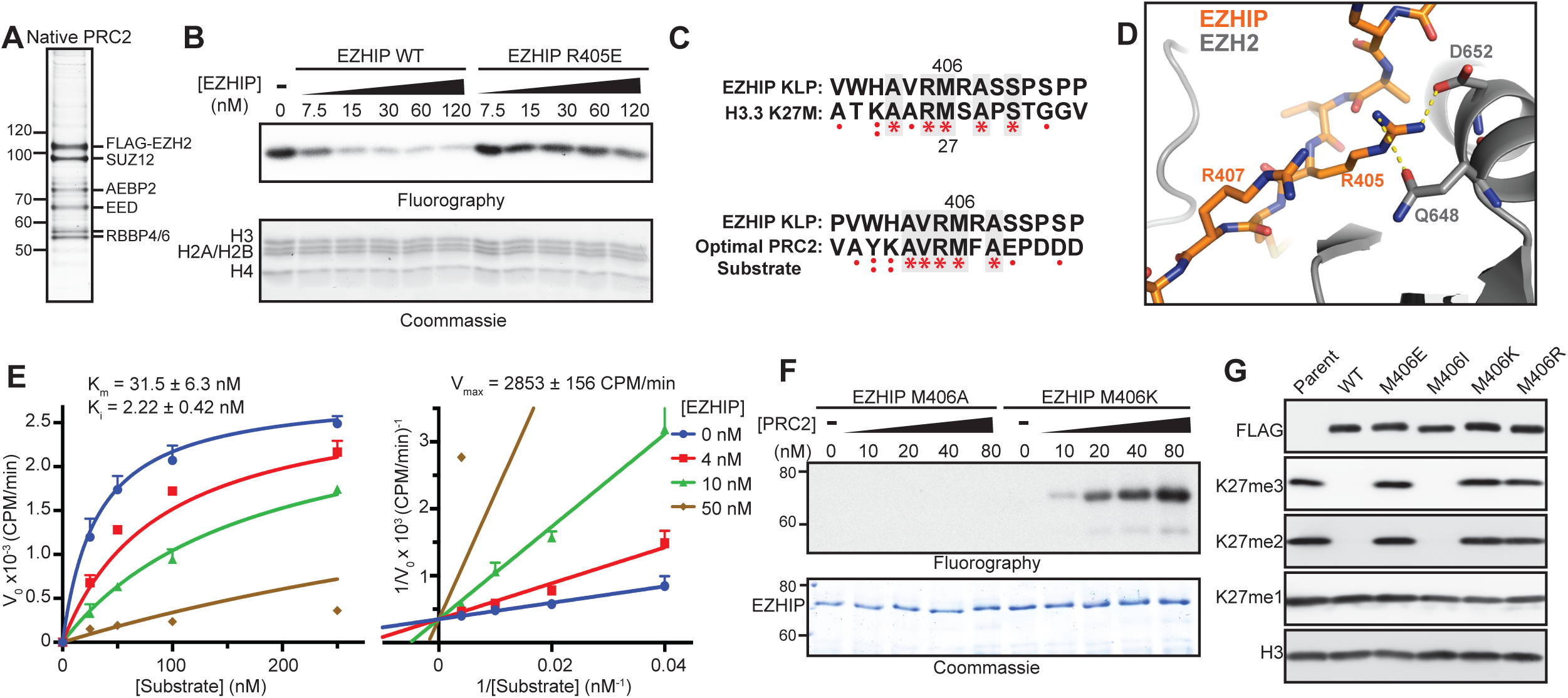
EZHIP is a competitive inhibitor of PRC2. (A) Silver stained SDS-PAGE gel showing the components of native PRC2 complex purified from HeLa cells. (B) *In vitro* methyltransferase reactions with PRC2 and oligonucleosome substrate. Full length recombinant EZHIP WT or R405E mutant purified from *E. coli* was titrated into the reaction mixture as shown. Half of the reaction was subjected to SDS-PAGE followed by fluorography and the other half was used for quantification by scintillation counting (Figure S2A). (C) Sequence alignment showing the similarity between lysine(27)-to-methionine mutated optimal PRC2 substrate and K27M-like EZHIP peptide (top), and H3K27M and K27M-like EZHIP peptide (bottom). (D) Formation of salt bridges between EZHIP R405 (or H3 R24) with EZH2 D652 (2.4Å) and Q648 (3.5Å), while R407 residue is exposed to solvent. (E) Increasing concentrations of oligonucleosome substrates were incubated with PRC2, SAM and varying concentrations of EZHIP inhibitor for 6 min. K_i_ of inhibition was determined by fitting Michaelis-Menton and Lineweaver-Burk curves demonstrating a competitive mode of inhibition. (F) 0.3 µM EZHIP M406A or M406K mutant proteins were incubated with increasing concentrations of PRC2 with 1 µM ^3^H-SAM and 25 µM H3K27me3 peptide for 1 hr. Reaction was subjected to SDS-PAGE followed by fluorography. (G) Immunoblots of 293T overexpressing EZHIP WT or M406E/I/K/R mutants.

Interestingly, the absolutely conserved twelve residue sequence in the C-terminus of EZHIP is remarkably similar to the sequence surrounding lysine 27 in the histone H3 K27M peptide (Figure 2C). With the exception of methionine 406, the EZHIP conserved peptide is remarkably similar to the calculated ‘optimal’ substrate sequence previously defined for PRC2 ^32^. PRC2 has a noteworthy preference for an arginine at −1 position relative to the substrate lysine, while the identity of the residue at the +1 position is seemingly less critical for optimal activity ^32^. We were able to model side chains of residues in the conserved EZHIP peptide on the H3K27M peptide observed in the PRC2-H3 K27M co-crystal structure without major steric hindrance ^12^ (Figure S2B). Interestingly, the critical R405 residue in EZHIP likely favorably interacts with D652 and Q648 of EZH2, whereas R407 was exposed to the solvent (Figure 2D, S2C). These observations provide a structural and biochemical basis for our finding that R405E, but not R407E, abrogated PRC2 inhibition by EZHIP (Figure 1J, 2B) and suggest that the EZHIP conserved sequence, henceforth referred to as <u>K</u>27M-<u>L</u>ike <u>P</u>eptide (**KLP**), makes direct contacts with EHZ2 active site in a similar manner to the H3 K27M peptide.

The strong inhibition of PRC2 methyltransferase activity led us to determine the detailed mechanism of action and Michaelis-Menten kinetic parameters associated with nucleosome substrates as a function of EZHIP concentration. We found that EZHIP inhibition appears competitive on nucleosomes (K_i_ = 2.22 ± 4.2 nM), indicated by a linear dependence of the IC_50_ values on the concentration of substrate (Figure 2E). This inhibition constant of 2.22 ±0.4 nM is comparable to K_i_ of inhibition by H3 K27M nucleosomes (2.1 ±0.9 nM),^14^ under similar reaction conditions.

The competitive mode of inhibition and the notable similarity between the KLP sequence to K27M and PRC2 substrates led us to hypothesize that EZHIP interacts directly with the active site of EZH2. Therefore, we reasoned that substituting a lysine for M406 would transform EZHIP into a PRC2 substrate. Indeed, we find that the full-length recombinant EZHIP M406K is a remarkably good substrate for PRC2, while the full-length M406A protein is not methylated by PRC2 to any measurable amount (Figure 2F, S2E). These data lead us to conclude that M406 binds to the active site of EZH2, likely through van der Waals interactions with a quartet of highly conserved aromatic residues lining the active site, and inhibits PRC2 through a K27M-like mechanism ^12-14,33,34^.

Previously, we found that histone H3 transgenes containing methionine or isoleucine at position 27 were capable of inhibiting PRC2 *in vitro* and decreasing H3K27me3 when expressed ectopically in cultured cells ^13^. Similarly, we found that an *EZHIP* transgene containing isoleucine at M406, but not acidic (glutamic acid) or basic (lysine and arginine) amino acid substitutions, reduced H3K27me2/3 in cells to a similar extent as wildtype *EZHIP* (Figure 2G). These data further confirmed our conclusion that EZHIP conserved peptide interacts with the aromatic cage residues of EZH2 active site.

### EZHIP KLP is sufficient and necessary to inhibit PRC2

Our *in vitro* and *in vivo* data indicate that a K27M-like peptide (KLP) in the C-terminus of EZHIP is necessary to inhibit PRC2 catalytic activity. We next sought to determine if the KLP was sufficient to inhibit PRC2 activity *in vitro*. To this end, we assessed recombinant PRC2 activity on histone H3 (18-37) peptide substrates in the presence of H3 K27M (18-37) or KLP (403-423) peptides (Figure 3A, B). In this side-by-side comparison, we found that the KLP was a more potent inhibitor of PRC2 activity (IC_50_= 4.1 µM) than the H3 K27M peptide (IC_50_ = 27.87 µM). Consistent with the critical role of M406 for KLP function, a M406E peptide did not inhibit PRC2 activity *in vitro* (Figure S1A). Additionally, we found that the IC_50_ correlated positively with the peptide substrate concentration (Figure 3D, S3B), a finding consistent with the competitive mode of inhibition that we found for full-length EZHIP and nucleosomal substrates (Figure 2D, E). In addition to EZH2, mammals contain a second Enhancer of Zeste (Ez) homolog called EZH1 that complexes with PRC2 subunits and methylates H3K27 ^35,36^. EZH1 peptides were identified in the EZHIP co-immunoprecipitated material that was analyzed by mass spectrometry (Figure S1D). We found that the KLP (IC_50_= 11.27 µM) and K27M (IC_50_= NA) inhibited EZH1-containing recombinant PRC2 *in vitro* (Figure 3A, C).

**Figure 3:**
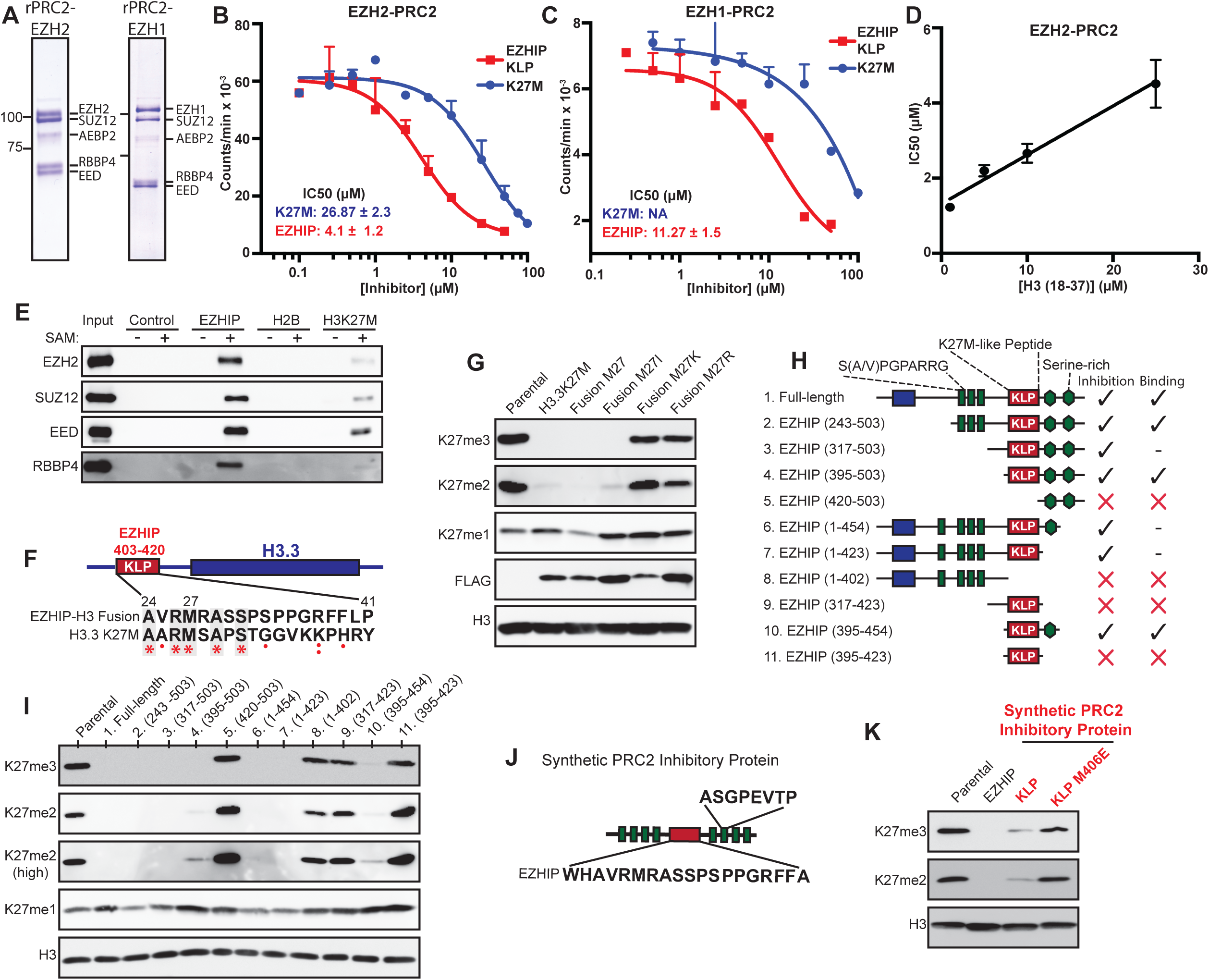
K27M-like conserved sequence in EZHIP is necessary and sufficient to inhibit PRC2 activity. (A) Coomassie stained SDS-PAGE gel displaying the components of recombinant PRC2 purified from SF9 cells. (B and C) *In vitro* methyltransferase reactions with rPRC2-Ezh2 (B) or rPRC2-Ezh1 (C) and peptide substrates with increasing concentrations of EZHIP (403-423) or H3K27M (18-37) peptides. Variable slope, four parameter Hill curve was fitted to determine the IC_50_ of PRC2 inhibition under these conditions. (D) IC_50_ of PRC2 inhibition by EZHIP peptide at different substrate (H3 (18-37)) concentrations (Fig S3A) is plotted against the corresponding substrate concentration. A linear positive correlation between IC_50_ and [substrate] is consistent with the competitive mode of inhibition. (E) Streptavidin agarose beads bound to biotinylated EZHIP (403-422), H2B (1-21) or H3K27M (18-37) peptides were incubated with 3 µg of recombinant PRC2 in the presence or absence of 40 µM SAM. After washing 3x, eluted samples were subjected to immunoblotting. (F and G) Immunoblots of whole cell extracts of 293T overexpressing HA-FLAG-tagged H3-EZHIP fusion protein with M27 or M27I/K/R mutants as shown in F. (H and I) Immunoblots of 293T cells overexpressing truncated EZHIP protein as shown in H. Blue, red, and green boxes represent the hotspot for ependymoma-associated mutations, K27M-like peptide (KLP), and N-terminal short tandem repeats of EZHIP respectively, green hexagons represent serine-rich regions. (J and K) Immunoblots from lysates prepared from 293T cells overexpressing synthetic protein as shown in J. Green and red boxes represent short tandem repeats and KLP or K27M sequences respectively.

Our previous studies revealed that methionine in the H3 K9M peptide occupies the active site of its cognate methyltransferase enzyme, G9a and forms a high-affinity ternary complex only in the presence of the co-substrate S-adenosyl methionine (SAM) ^33^. Consistent with the formation of a ternary complex, kinetic analysis indicated that H3 K9M is an uncompetitive inhibitor of G9a with respect to SAM. We proposed that a SAM-induced conformational change allows the active site of the methyltransferases to stably interact with K-to-M histones found in human cancers. A similar SAM-dependent interaction between H3 K27M and PRC2 has also been described ^12^. In addition to H3 K27M, we found that the KLP interacted with PRC2 in a SAM-dependent manner (Figure 3E) and that this association was abolished by the M406E substitution (Figure S3C). The KLP-PRC2 interaction was more stable than the K27M-PRC2 interaction at high NaCl concentrations. Together, these data suggest that a high affinity, SAM-dependent ternary complex forms between SAM-bound PRC2 and EZHIP (Figure S3D). Consistent with our finding that the KLP interacts with the active site of EZH2, preincubation of PRC2 with H3 K27M and SAM abrogated PRC2 association with the KLP (Figure S3E).

Next, we synthesized a histone H3-EZHIP fusion protein to assess if the KLP, alone, in the context of the histone H3 tail could lower H3K27me2/3 levels in cells. We replaced residues 24-41 of histone H3 with EZHIP residues 403-420 spanning the KLP sequence. H3-EZHIP fusion transgenes containing methionine or isoleucine at residue 27 lowered global H3K27me2/3 levels, while substitution of lysine or arginine had no effect (Figure 3G, S1B). Taken together, these data suggest that the K27M-like sequence in EZHIP is necessary and sufficient for PRC2 inhibition *in vivo*.

In our *in vitro* PRC2 assays, we observed that full-length EZHIP protein inhibits PRC2 at a lower concentration compared to KLP itself, suggesting the presence of a secondary PRC2 binding domain (Figure S3F). EZHIP is predicted to lack any secondary structure and KLP is flanked by intrinsically disordered regions (IDRs) containing short-tandem repeats, which are common in IDRs involved in protein-protein interactions (citation). We hypothesized that IDRs flanking KLP may facilitate EZHIP-PRC2 interaction and inhibition. To identify secondary PRC2 binding site within EZHIP, we generated a series of *EZHIP* deletion transgenes and assessed their potential to inhibit and bind PRC2 *in vivo* (Figure 3H-I, S3G). We found that deletion of C-terminal or N-terminal residues from EZHIP (fragments 2, 3, 4, 6, 7) lowered H3K27me2/3 comparable to full-length EZHIP. An additional deletion of KLP from these transgenes (fragments 5, 8) abrogated its ability to inhibit and bind PRC2, underscoring the necessity of KLP for its activity. We observed that KLP by itself (fragments 9, 11) was unable to reduce H3K27me2/3 *in vivo*, mirroring its lower inhibitory potential *in vitro* compared to full-length protein (Figure S3F). Interestingly, addition of either the C-terminal serine-rich IDRs or N-terminal short tandem repeats (fragments 4, 7, 10) significantly enhanced KLP binding and inhibition of PRC2. Since the flanking IDRs were unable to reduce H3K27me2/3 by themselves (fragments 5, 8) and were individually dispensable, we conclude that KLP is the primary PRC2 recognition and inhibitory domain within EZHIP and flanking IDRs enhance its inhibitory potential by making weak, non-specific interactions with PRC2.

To directly test the hypothesis that KLP is sufficient to lower H3K27me2/3 and utilizes flanking IDRs for strengthening EZHIP-PRC2 interaction, we synthesized an artificial protein containing KLP flanked by four intrinsically-disordered short, tandem repeats on each side (Figure 3J). We found that KLP sequence, but not M406E, embedded within flanking IDRs reduced H3K27me2/3 *in vivo* (Figure 3K). These data further support our conclusion that EZHIP KLP is sufficient to inhibit PRC2 *in vivo*.

### *EZHIP* expression leads to K27M-like genomic distribution of H3K27me

Expression of *EZHIP* transgenes caused a marked reduction in H3K27me2/3 in various cell types through inhibition of PRC2 activity in a K27M-like mechanism. We used chromatin immunoprecipitation followed by DNA sequencing (ChIP-Seq) and RNA sequencing to profile changes in gene expression and the chromatin landscape in MEFs expressing *EZHIP*. First, we generated MEF cell lines from E13.5 mouse embryo that contain loxP sites flanking exons 3-6 of the *EED* gene ^37^. We used these MEF lines to derive isogenic transgenic H3.3 K27M and *EZHIP* cell lines to directly compare chromatin and gene expression changes caused by these two PRC2 inhibitor proteins. Cre-mediated excision of *EED* served as a control for chromatin and gene expression changes caused by genetic depletion of PRC2 in our MEF cell line.

Consistent with our mass spectrometry and immunoblot data, we observed a genome-wide reduction of H3K27me3 in cells expressing H3.3 K27M or *EZHIP* as measured by the average normalized read counts of the immunoprecipitated chromatin (Figure S5A, 4C). Additionally, *EZHIP* or H3.3 K27M expressing cells exhibited an 80% decrease in the total number of H3K27me3 peaks (Figure S4A, B). Though some H3K27me3 peaks disappeared completely in *EZHIP* and H3.3 K27M cells, we noted a striking difference in distribution of H3K27me3 among the peaks that remained; while H3K27me3 is usually found in broad domains in wildtype cells, we noted that the peak width decreased substantially in cells expressing *EZHIP* or H3.3 K27M (Figure 4A, B). Similar changes in H3K27me3 ChIP-seq profiles have been observed in DIPG cells containing the H3 K27M mutant histone ^8,15-19^.

**Figure 4:**
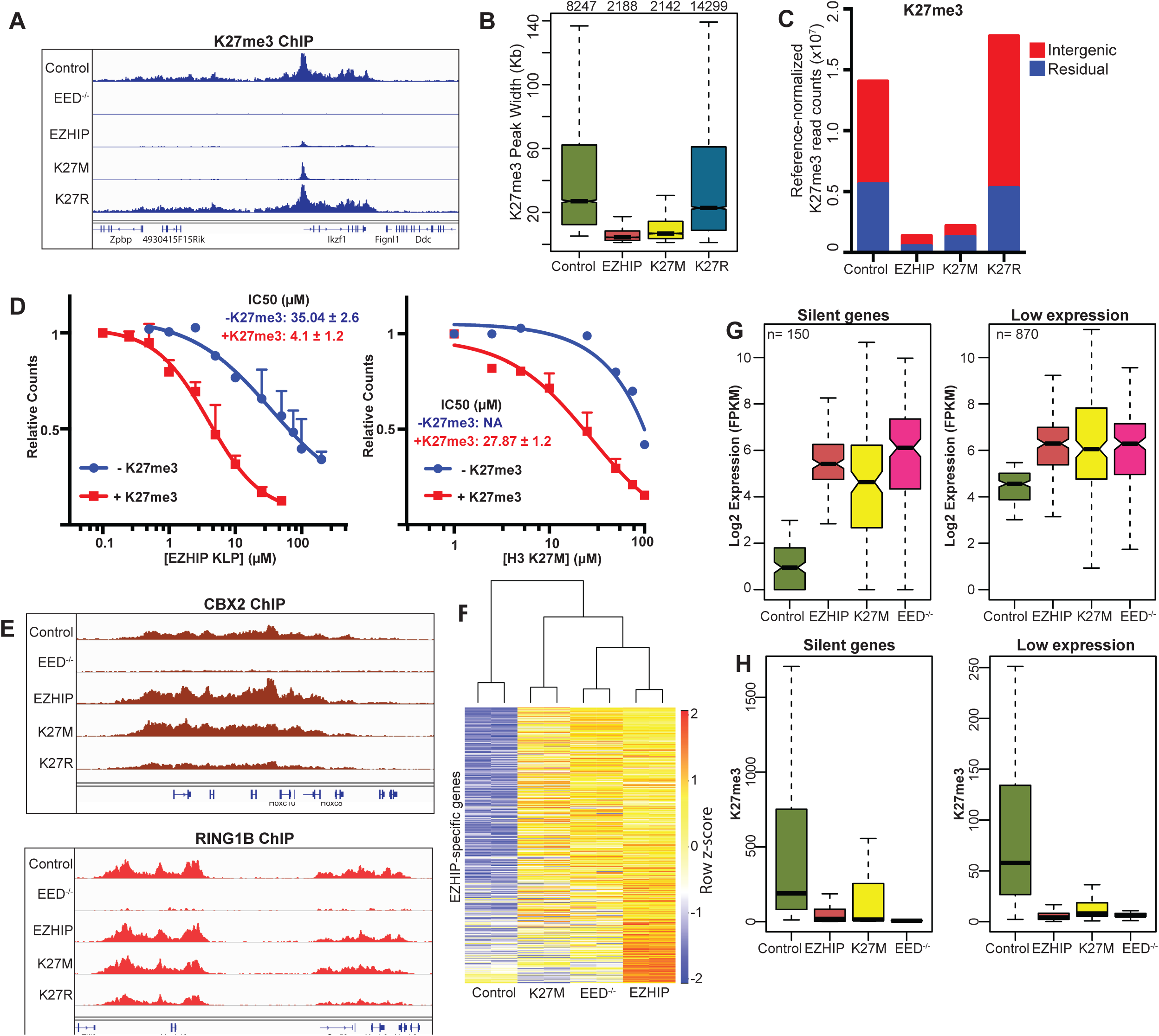
EZHIP leads to global loss of H3K27me3 and a concurrent upregulation of silenced genes. (A) Genome browser representation of reference-normalized H3K27me3 ChIP-Seq profile in mouse embryonic fibroblasts overexpressing H3K27M, H3K27R, EZHIP or control cells in a 500 kb region of the genome. (B) Boxplot showing the genomic region occupied by H3K27me3 peaks in the indicated samples. (C) The number of reference normalized H3K27me3 read densities associated with intergenic or residual H3K27me3 enriched regions are displayed as barplot. (D) *In vitro* PRC2 reactions using H3 18-37 peptide in the absence or presence of 20 µM H3K27me3 stimulatory peptide, with increasing concentrations of EZHIP KLP (left) or H3 K27M (right). Variable slope, four parameter Hill curve was fitted to the data to determine IC_50_ values. (E) Genome browser representation of RPKM normalized Cbx2 and Ring1b ChIP-Seq profiles from MEFs overexpressing H3K27M, H3K27R, EZHIP or control cells. (F) Heatmap displaying the expression pattern of genes upregulated in cells overexpressing EZHIP. Unguided hierarchical clustering was used to generate dendrogram shown above the heatmap. (G) Boxplot displaying the expression of silent (left) and low-expression (right) genes. (H) Boxplot displaying the reference normalized K27me3 enrichment at the promoters (5kb upstream of TSS) of genes displayed in G.

Previous studies demonstrated that PRC2 is allosterically activated via interaction of the EED subunit with either H3K27me3 or JARID2-K116me3 ^12,38,39^. These structural and biochemical studies found that the SET domain of EZH2 adopts a catalytically productive conformation when the EED subunit is engaged with the trimethylated H3K27 or JARID2-K116. Allosteric stimulation of PRC2 allows it to catalyze H3K27me3 on the neighboring nucleosomes *in cis* after initial recruitment to CpG islands, giving rise to broad H3K27me3 domains that are observed *in vivo*. Our ChIP-Seq data showed that *EZHIP* expression led to loss of H3K27me3 from secondary spread sites while retaining it at the recruitment sites. This loss of spreading led us to hypothesize that the allosterically activated form of PRC2 is more sensitive to inhibition by EZHIP. Indeed, we found that the H3.3 K27M (+K27me3 IC_50_ = 27.87 µM, compared to -K27me3 IC_50_ = NA) and KLP (+K27me3 IC_50_ = 4.1 µM, compared to -K27me3 IC_50_ = 35.04 µM) inhibited allosterically activated PRC2 at a lower concentration *in vitro* (Figure 4D). A similar mechanism has been proposed for H3 K27M oncohistone where H3K27me3-bound PRC2 interacts with the H3 K27M peptide with higher affinity ^12^.

The genome-wide reduction of H3K27me3 led us to investigate whether the Polycomb Repressive Complex 1 (PRC1) distribution was altered by *EZHIP* or H3.3 K27M expression. Despite the overall decrease in H3K27me3 peak number and peak width, we found that the number and general distribution of PRC1 subunits CBX2 and RING1B remained unchanged by the expression of *EZHIP* or H3.3 K27M (Figure 4E, S5C-D). This finding suggests that the residual, H3K27me3-modified nucleosomes present in H3.3 K27M and *EZHIP*-expressing cells retain the capacity to recruit PRC1 complex.

We performed total RNA-sequencing to identify EZHIP-and H3 K27M-mediated changes in gene expression. Consistent with a loss of H3K27me3, we found up-regulation of ∼500 genes in cells expressing *EZHIP*. Importantly, expression of H3 K27M or genetic depletion of PRC2 led to similar upregulation of these genes (Figure 4F-G). Specifically, we observed upregulation of silenced and lowly expressed genes that contains high levels of H3K27me3 in their promoters (Figure 4G-H, S5E-F). Upregulation of these genes correlated with the loss of H3K27me3 from their promoters (Figure 4G-H). Together, our data show that expression of *EZHIP* or H3.3 K27M in an isogenic, cell culture system leads to a chromatin and gene expression profile that largely reflects loss of PRC2-mediated gene silencing.

### Ependymomas expressing EZHIP display aberrant silencing of CDKN2A by increasing local H3K27me3

Our *in vivo* experiments indicate that EZHIP and H3 K27M generate similar genome-wide chromatin and gene expression profiles. We next analyzed H3K27me3 ChIP and RNA sequencing data from supratentorial (*EZHIP* negative, high H3K27me3) and PFA ependymomas (*EZHIP* positive, low H3K27me3). Our H3K27me3 ChIP sequencing analyses show that PFA ependymomas exhibit a similar chromatin profile to our MEFs expressing *EZHIP*; reduced peak width at all H3K27me3 peaks and a reduced number of total peaks (Figure 5A, B). We analyzed K27M-containing DIPG ChIP sequencing data and found a remarkable overlap in the location of residual H3K27me3 peaks found in K27M-containing DIPGs and PFA ependymomas (Figure 5C). These data support previously noted clinical and biological similarities between PFA ependymomas and DIPG tumors ^21,23,40^.

**Figure 5:**
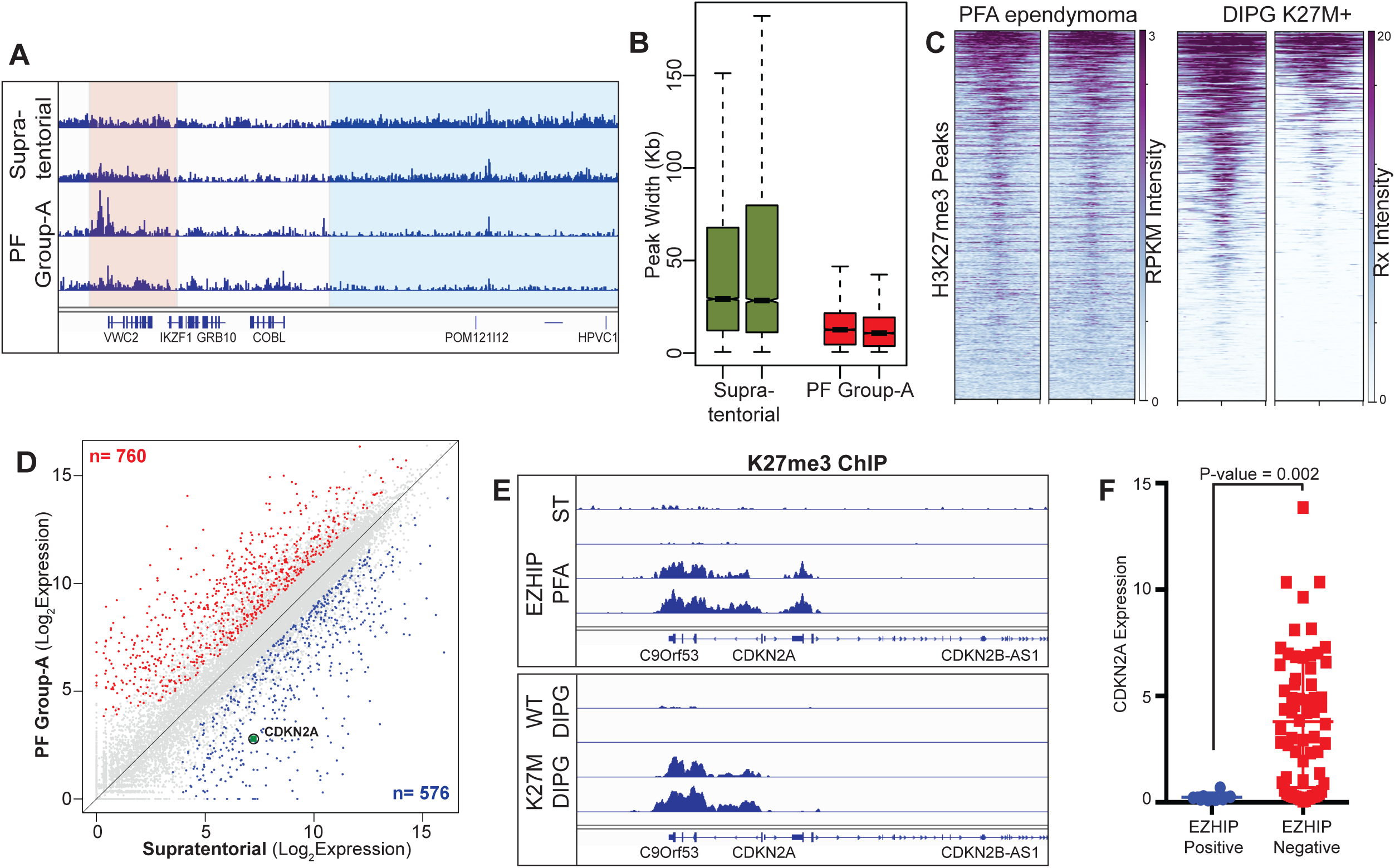
PFA tumors expressing EZHIP display lowered expression of CDKN2A by increasing “local” K27me3. (A) Genome browser representation of RPKM normalized K27me3 ChIP-Seq profile in Posterior Fossa Group-A ependymomas overexpressing EZHIP and supratentorial ependymomas. Blue and red boxes represent intergenic and residual (retained) K27me3 respectively. (B) Boxplot displaying the genomic region occupied by K27me3 peaks in supratentorial and PFA ependymomas. (C) Heatmap displaying the normalized K27me3 enrichment at peaks retained in PFA ependymomas overexpressing EZHIP in PFA (left) and DIPG cell lines containing H3K27M mutations (right). (D) Scatter plot displaying the expression of all genes in PFA and supratentorial ependymomas. Red and blue points represent up-and down-regulated genes in PFA tumors relative to supratentorial tumors. Encircled, green point represents the expression of CDKN2A gene. (E) Genome browser representation of K27me3 ChIP-Seq profile at the CDKN2A locus in PFA and supratentorial tumors (top); and H3.3WT or H3.3K27M DIPG lines (bottom). (F) Expression of CDKN2A gene in ependymomas with low (blue, n = 7) or high (red, n = 71) expression of EZHIP, as measured by FPKM values. Ependymomas were grouped based on their expression of EZHIP. P-value was calculated using non-parametric t-test.

We plotted the log2 expression profile of genes in PFA and supratentorial ependymomas to identify gene expression changes possibly caused by EZHIP expression (Figure 5D, S5A) Using STRING analysis, we found enrichment of several gene ontology terms related to organism development and cell differentiation (Figure S5B). Earlier studies found that K27M-containing DIPGs exhibit global reduction in H3K27me3 and retention of H3K27me3 peaks at many genomic loci ^8,15-19^. The *CDKN2A* tumor suppressor gene is one locus that is silenced in a H3K27me3-dependent manner in K27M-containing DIPG cell lines and in H3.3 K27M glioma model systems ^18,41,42^. Similarly, we found a 20-fold down-regulation of *CDKN2A* expression in PFA ependymomas. ChIP sequencing tracks show high levels of H3K27me3 at the *CDKN2A* locus in K27M-containing DIPG and EZHIP-expressing PFA ependymomas (Figure 5E). Additionally, ependymomas containing high *EZHIP* expression exhibit reduced levels of *CDKN2A* as compared to ependymomas with low EZHIP (Figure 5F). These results suggest that retention of PRC2 activity at *CDKN2A* in ependymomas expressing *EZHIP* may be a major contribution to tumorigenesis. Overall, these data support a model whereby chromatin-based silencing of tumor suppressors by mutation (K27M) or aberrant gene expression (*EZHIP*) facilitates formation two hindbrain tumor types (DIPG and PFA ependymomas).

## DISCUSSION

In this study we demonstrated that expression of *EZHIP*, a putative driver of PFA ependymomas, lowers global H3K27me3 levels through inhibition of PRC2 methyltransferase activity. Our biochemical experiments show a direct and robust interaction with the core-PRC2 subunits. Moreover, the findings from our *in vitro* studies of EZHIP parallel results obtained from studies of the H3 K27M oncohistone found in DIPG and other high-grade midline gliomas. Both EZHIP and K27M contain short sequences that are potent competitive inhibitors of PRC2 in regard to the nucleosome substrate, interact with PRC2 in a SAM-dependent manner, and contain a critical methionine necessary for inhibition of PRC2 methyltransferase activity.

*EZHIP* homologs are found exclusively in placental mammals. Its expression pattern in normal development appears to be tightly restricted to spermatogonia, suggesting that the protein likely plays a role in spermatogenesis ^43^. Genes involved in sexual reproduction have been noted to evolve rapidly, especially in rodents ^44^. Interestingly, we find that *EZHIP* homolog sequences exhibit the most divergence among rodent species (**Figure 1H**). *EZHIP* may have evolved to dynamically regulate PRC2 activity during spermatogenesis in mammals, and future studies will address questions related to the regulation of H3K27 methylation during spermatozoa development.

Missense mutations in *EZHIP* are reported in fewer than 10% of PFAs ^26^. We found that missense mutations do not affect the H3K27 methylation lowering activity of EZHIP *in vivo*, suggesting that these tumor mutations do not alter the EZHIP protein function. Notably, nonsense or frameshift mutations that would alter the expression of the inhibitory EZHIP C-terminus have not been reported in PFA tumors. Instead, we hypothesize that the mutations promote expression of *EZHIP* through altering one or more *cis*-acting regulatory elements found within the gene body. Enhancers and other *cis*-acting genetic elements frequently occur within introns of genes. The *EZHIP* gene is, in all placental mammals, a long contiguous gene that lacks introns, thus functional mutations to intragenic regulatory elements will inadvertently alter the protein sequence. It is likely that PFA tumors that lack *EZHIP* mutations drive expression of the gene through other genetic or epigenetic mechanisms.

In addition to PFA ependymomas, *EZHIP* expression is implicated in uterine neoplasms, as a *MBTD1-CXORF67* fusion gene was reported to occur in several endometrial stromal sarcomas ^45^. Importantly, the MBTD1-CXORF67 fusion protein contains the C-terminus of EZHIP and the highly conserved K27M-like sequence, suggesting that PRC2 inhibition and loss of H3K27 methylation may support tumorigenesis of this subtype of endometrial cancer. In support of this model, other fusion proteins between PRC2 subunits and the zinc finger-containing JAZF1 protein have been reported; JAZF1-PHF1 fusion proteins occur in a small number of endometrial stromal tumors, while a JAZF1-SUZ12 fusion protein reportedly occur in 75% of this tumor type that accounts for less than 10% of all uterine tumors ^46-49^. The JAZF1-SUZ12 protein is reported to dysregulate PRC2 activity and lower H3K27me3 levels in cells ^46^.

Curiously, no loss-of-function mutations of the core PRC2 subunits are reported to occur in pediatric DIPG or PFA ependymomas. On the contrary, multiple inactivating alleles of the PRC2 core subunits (*EED* and *SUZ12*) are reported in >80% of malignant peripheral nerve sheath tumors (MPNSTs) ^50^. Based on these data, we previously hypothesized that some PcG-mediated gene repression remains intact in cells expressing the K27M mutant histones, and that PRC2 activity is required for K27M-mediated transformation ^34^. In support of residual PRC2 activity in cells, previously studies have found residual H3K27me3 in DIPG cells. Additionally, ChIP seq data from our isogenic cell lines revealed that expression of H3 K27M or EZHIP led to a reduction in total H3K27me3 peak numbers and peak width. The H3K27me3 nucleosomes retained at sites of PRC2 recruitment are functionally critical for DIPG growth as these cells are sensitive to pharmacological inhibitors of PRC2 ^18,19^. We speculate that PFA-ependymomas that express high levels of EZHIP will also exhibit differential sensitivity to EZH2 inhibitors.

Previously, we and others have proposed that inhibition of PRC2 by H3 K27M prevents spreading of H3K27me3 into the secondary sites and intergenic regions, but allows its residual activity at primary, strong PRC2 sites at CpG islands ^16,18,34^. This residual PRC2 activity is evident from an apparent local retention of residual H3K27me3 at specific loci. It is known that certain PTMs can positively or negatively regulate PRC2 methyltransferase activity, and the enzyme is stimulated by its own reaction product, namely H3K27me3 through interaction with the EED subunit ^38^. In addition to serving as a possible recruitment mechanism, this feedforward mechanism is hypothesized to aid in establishing positive reinforcing loops that may facilitate spreading of H3K27me3 and Polycomb Repressive Complexes *in cis*. In contrast, the presence of the active chromatin marks, H3K4me3 and H3K36me3, each lead to a diminution in PRC2 activity ^51-53^. These studies and others suggest that PRC2 is a dynamic signal integration device capable, through accessory subunits, of converting various inputs from the local chromatin context into an appropriate enzymatic output that ultimately allows controlled spreading of the polycomb transcriptionally silenced state.

In this study, we have observed that EZHIP has remarkably better inhibitory activity towards PRC2 that is allosterically activated by H3K27me3 (>8-fold difference in IC_50_ values). In the presence of high levels of H3K27me3, EZHIP efficiently inhibits stimulated-PRC2, leading to loss PRC2 spreading and an eventual recession of H3K27me3 back to sites of PRC2 recruitment. We speculate that EZHIP inhibitory activity directed towards PRC2 will diminish as H3K27me3 levels recede, allowing PRC2 to once again catalyze H3K27me3 on nucleosomes proximal to sites of recruitment. Hence, we propose EZHIP or K27M inhibition of PRC2, and its activity on nucleosomes are in a state of equilibrium that results in an optimal level of localized H3K27me3 conducive for PFA or DIPG development (Figure 6).

**Figure 6:**
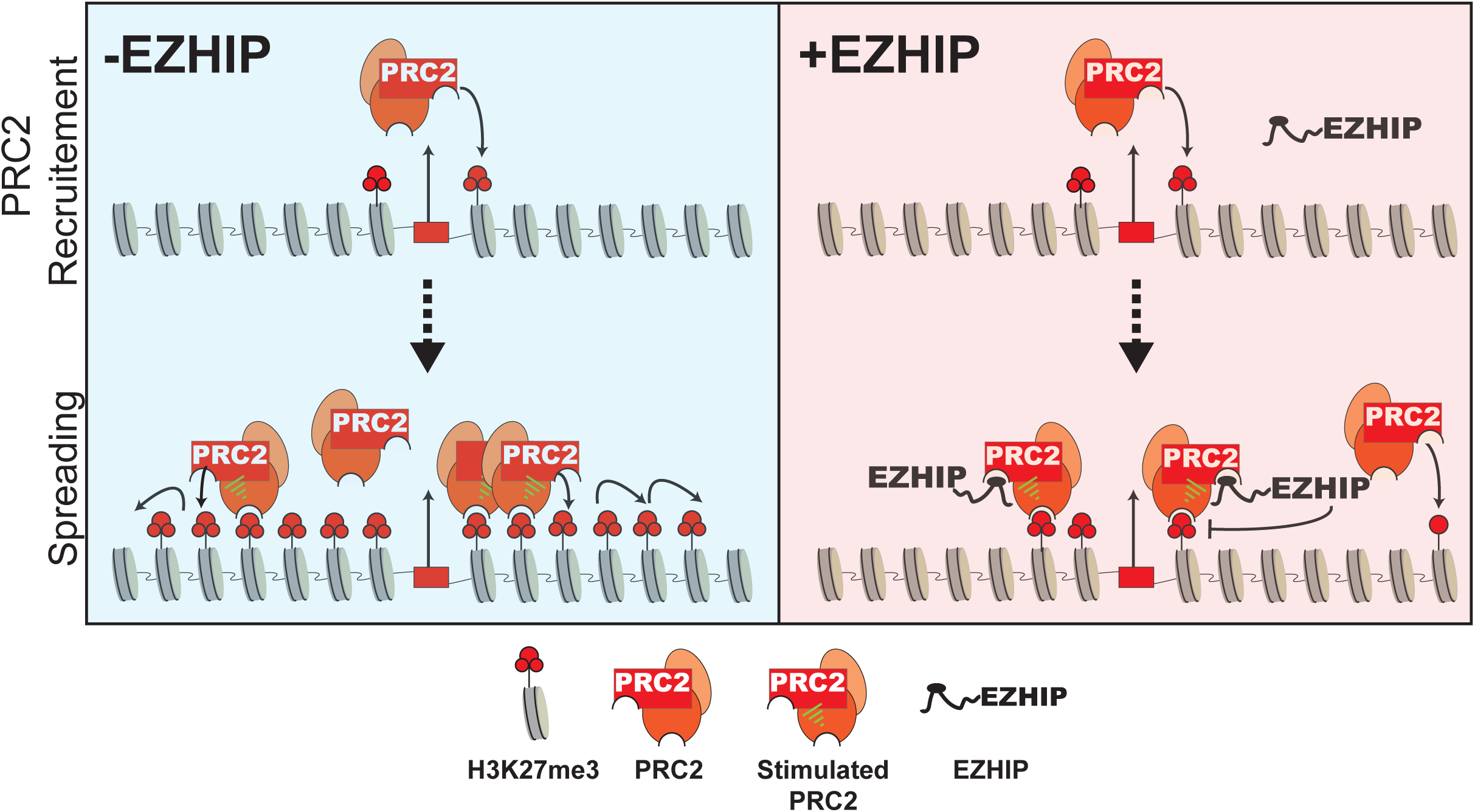
Oncohistone-mimic, EZHIP blocks H3K27me3 spreading by inhibiting allosterically-stimulated PRC2. Schematic depicting the molecular mechanism by which EZHIP expression leads to the loss of H3K27me3 spreading. In cells lacking EZHIP expression (in blue), PRC2 is recruited to CpG islands and catalyzes H3K27me3 at recruitment sites. H3K27me3-marked nucleosomes proximal to the recruitment site allosterically activate PRC2 and promote spreading *in cis* to form broad H3K27me3 domains. Similarly, in cells expressing EZHIP (red), PRC2 is recruited to CpG islands and can catalyze H3K27me3 at proximal nucleosomes due the weaker inhibition potential of EZHIP for unstimulated PRC2. However, spreading of H3K27me3 is blunted due to enhanced binding of EZHIP to allosterically stimulated PRC2. The formation of H3K27me3-PRC2-EZHIP ternary complex leads to inhibition of PRC2 spreading and provides a mechanism to explain the formation of narrow H3K27me3 peaks found in PFA ependymomas.

Cellular differentiation in normal development includes dynamic gain and loss of H3K27me3 at genes affecting cellular commitment. The expression of *EZHIP* or H3 K27M impairs the dynamic regulation of H3K27me3 by impeding its production by PRC2. The proposed cell-of-origin for PFA ependymomas is radial glia that serve as CNS progenitor cells, and a recent single cell sequencing study points to oligodendroglial progenitors as the likely cell-of-origin for DIPGs ^8,21,41,54,55^. Previous studies have found that H3 K27M impedes differentiation of neural precursor cells (NPCs) likely due to their inability to effectively silence genes involved in proliferation and multipotency ^18,41^. Additionally, these NPCs appear to adopt a more primitive stem state based on gene expression analysis. Here, we demonstrated that *CDKN2A* is silenced in *EZHIP*-expressing PFAs. *CDKN2A* encodes for critical regulators of the G1-S transition and is completely silenced in iPSCs and other stem cell populations ^56^. While the gene is eventually activated during lineage commitment, the aberrant silencing of *CDKN2A* is known to contribute to DIPG tumorigenesis ^18^. Future studies will address whether ectopic expression of *EZHIP* in radial glial cell populations lead to similar changes in gene expression and lineage commitment.

The sites of H3K27me3 found in PFA ependymomas and DIPG cells likely represent CpG islands that are the original recruitment sites for PRC2 in the cell-of-origin that gave rise to the tumor. We observed that a relatively small number of genes are dysregulated in *EZHIP* or H3 K27M expressing cells, despite the widespread reduction of H3K27me3 over many genes. We propose that many of these genes were not expressed due to redundant mechanisms of repression or by the inefficient transcription activation. We find that expression of H3 K27M or EZHIP lead to similar chromatin states and gene expression profiles in cells. While some PFA ependymomas contain the K27M mutation, *EZHIP* expression is the predominant mechanism for modulating PRC2 in these tumors. It is unknown whether DIPGs lacking the K27M mutation instead exhibit high levels of *EZHIP* expression. Additionally, it’s unclear why PFAs and DIPGs seemingly exhibit a preference for *EZHIP* and H3 K27M, respectively.

In summary, we demonstrate that EZHIP inhibits PRC2 activity in a K27M-like mechanism. We propose that aberrant expression of EZHIP contributes to PFA ependymoma tumorigenesis through dysregulation of PRC2-mediated gene repression. Defining the molecular pathways involved in mediating the oncogenic potential of EZHIP will identify therapeutic targets that may improve disease management and outcome.

## ACKNOWLEDGEMENTS

This research was supported by funding from P01CA196539 (to P.W.L., T.W.M., N.J., and B.A.G); the Greater Milwaukee Foundation (to P.W.L), the Sidney Kimmel Foundation (Kimmel Scholar Award to P.W.L.), a startup provided by the Wisconsin Institute for Discovery (to P.W.L.), and NIH grants; Innovator grant DP2OD007447 and R01GM110174 (to B.A.G.); Doris Duke Foundation Clinical Scientist Development Award (2016100), Sontag Foundation Distinguished Scientist Award (791165), Sidney Kimmel Foundation (444000) and K08CA181475 to S.V. A.B. is supported by Fonds de Recherche du Québec-Santé. This work was performed within the context of the I-CHANGE consortium and supported by funding from Genome Canada, Genome Quebec, The Institute for Cancer Research of the Canadian Institutes for Health Research (CIHR), McGill University and the Montreal Children’s Hospital Foundation. N.J. is a member of the Penny Cole lab and the recipient of a Chercheur Clinician Senior Award. We thank Dr. John Svaren for providing EED^f/f^ mice, and Drs. Rupa Sridharan and Coral Wille for helping with the isolation of mouse embryonic fibroblasts.

## Data Availability

The sequencing data reported in this paper are deposited at GEO database (GSE124839, GSE124743). Published data used in this paper were downloaded from GEO (GSE87779, GSE89452, and GSE118954). Raw mass spectrometry data reported in this paper have been deposited at ProteomeXchange (Accession ID: PXD013117).

## Supplementary Material

### Supplementary Figure Legend

**Figure S1:**
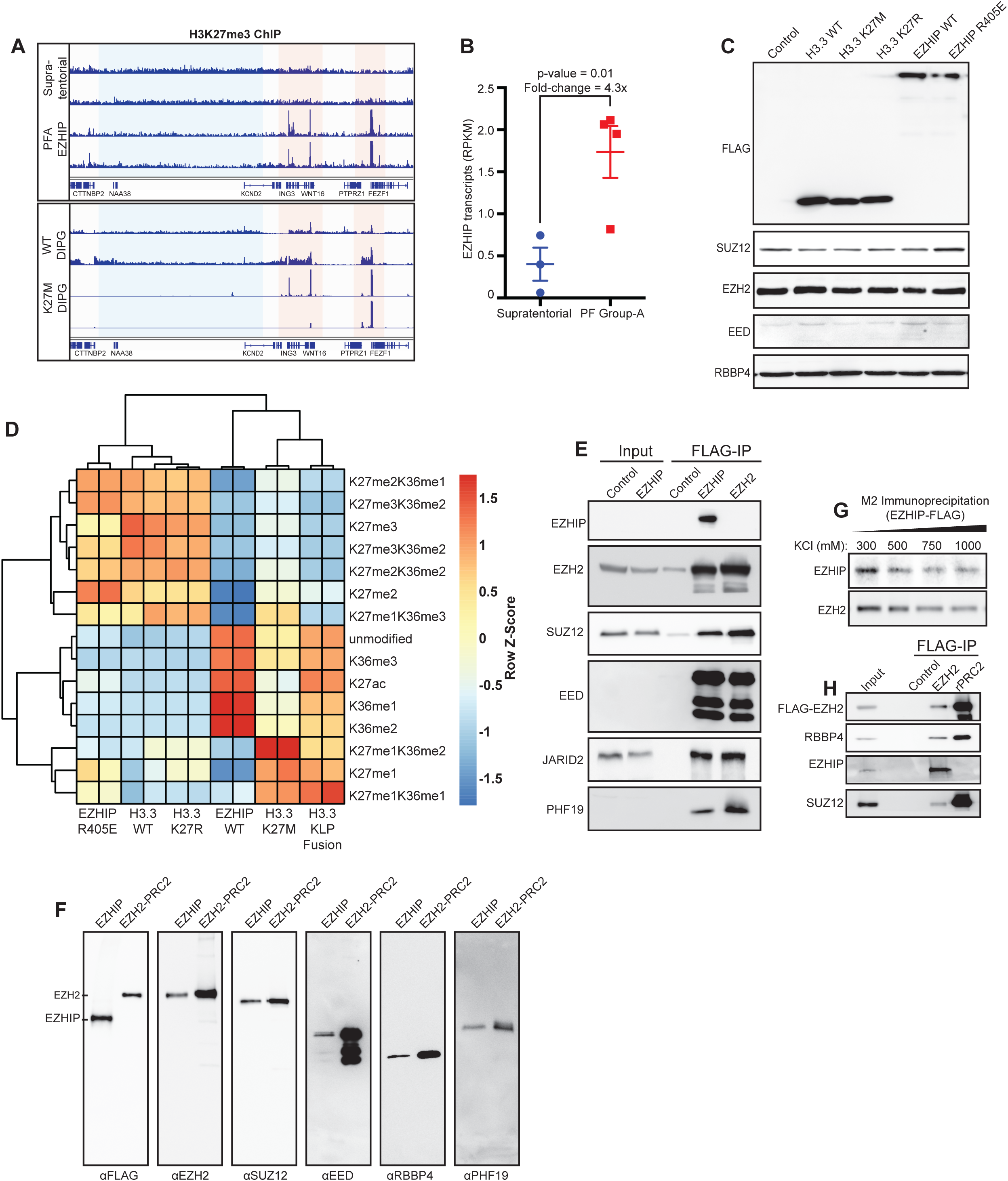
EZHIP forms a stable complex with PRC2 and lowers H3K27me3 *in vivo*. (A) Genome viewer representation of ChIP-Seq for H3K27me3 in a 6 Mbp region of the human genome. Red shaded region represents residual H3K27me3; broad, intergenic H3K27me3 in blue. (B) Plot displaying the number of EZHIP transcripts found in PFA and supratentorial ependymomas. Reads were RPKM normalized. (C) Immunoblots from whole cells lysates of 293T cells displaying the components of PRC2 complex overexpressing HA-FLAG-tagged H3.3 WT or K27M/R or FLAG-tagged EZHIP WT or R405E mutants. (D) Acid-extracted histones were subjected to bottom-up quantitative mass-spectrometry for measurement of histone PTM abundances. The relative abundance of H3K27, H3K36 and H3K9 modifications are plotted as heatmap. Fold change was determined as K27M/H3.3WT, K27R/H3.3WT, H3-EZHIP fusion/H3.3WT, EZHIPWT/EZHIP R405E. (E) Immunoblots of eluates from FLAG M2 immunoprecipitation from nuclear extracts of 293T cells overexpressing FLAG-tagged EZHIP or FLAG-tagged Ezh2. Nuclear extract from parental 293T cells was used as control. (F) M2 Immunoprecipitated material (EZHIP-FLAG or EZH2-FLAG) was subjected to anion-exchange chromatography. Immunoblots display the components of PRC2 in eluate corresponding to 350 mM KCl from anion exchange chromatography. (G) Immunoblots of eluates from FLAG M2 immunoprecipitation as in E, except with washes with buffers containing increasing concentrations of KCl. (H) Immunoblots of eluates from FLAG M2 immunoprecipitation from nuclear extracts of U2OS cells overexpressing FLAG-tagged EZH2. Nuclear extract was first incubated with unconjugated agarose beads, which was used as control.

**Figure S2:**
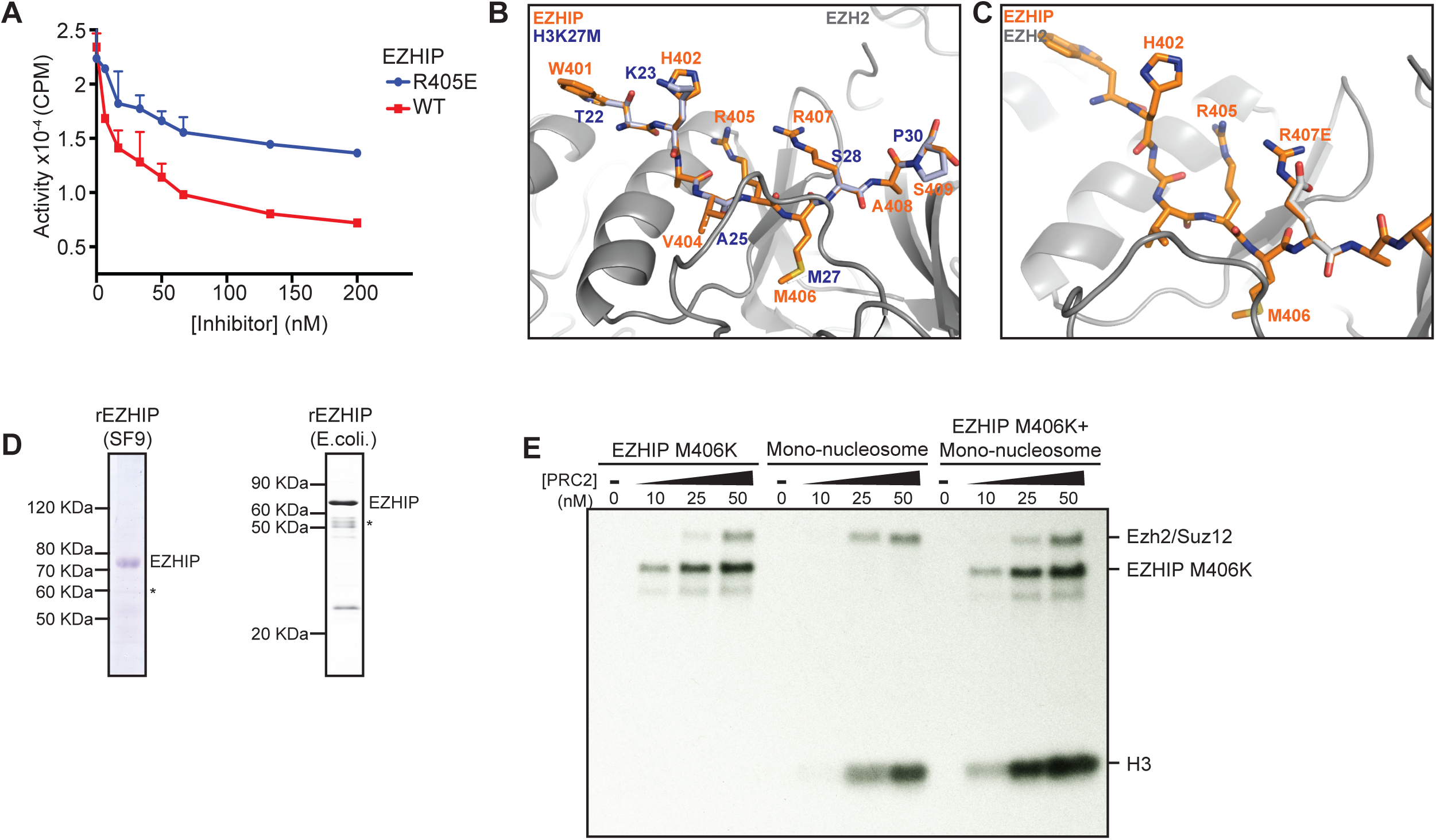
EZHIP conserved peptide mimics H3K27M peptide. (A) Quantification of *in vitro* methyltransferase reactions with PRC2 and oligonucleosome substrate. Full length recombinant EZHIP WT or R405E mutant purified from *E. coli* was titrated into the reaction mixture as shown. Half of the reaction was subjected to SDS-PAGE followed by fluorography (Figure 2B) and the other half was used for quantification by scintillation counting. (B) Side-chains of H3K27M peptide in the PRC2-H3K27M co-crystal structure (pdb: 5HYN) were mutated to model side chains of EZHIP conserved peptide. (C) Position of EZHIP R407 away from EZH2 surface, towards the solvent in the modelled co-crystal structure. (D) Coommassie stained SDS-PAGE gel showing recombinant EZHIP purified from SF9 cells and BL21 *E.coli*. (E) 0.3 µM EZHIP M406K mutant or recombinant nucleosomes or both were incubated with increasing concentrations of recombinant PRC2 with 1 µM ^3^H-SAM and 25 µM H3K27me3 peptide for 1 hr. Reaction was subjected to SDS-PAGE followed by fluorography.

**Figure S3:**
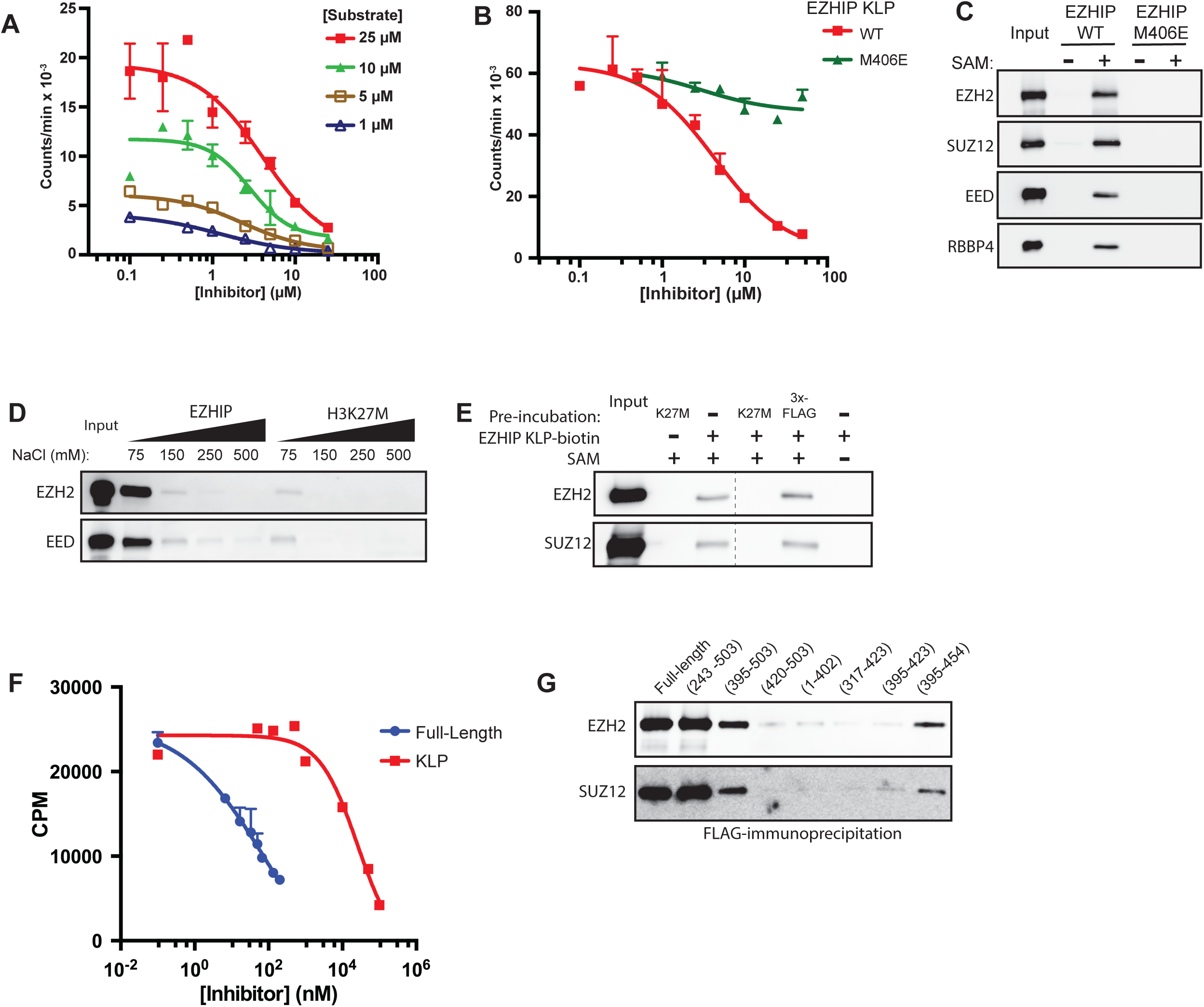
“K27M-like” EZHIP peptide binds and inhibits PRC2 complex. (A) *In vitro* PRC2 reactions using varying concentrations of H3 18-37 substrate peptides at increasing concentrations of K27M-like EZHIP 403-423 peptide. Variable slope, four parameter Hill curve was fitted to the data to determine IC50 values. (B) *In vitro* PRC2 reactions with rPRC2-Ezh2 and 50 µM peptide substrate with increasing concentrations of EZHIP WT or M406E peptides (403-423). (C) Peptide pulldowns were performed as described in Figure 3E with EZHIP WT or M406E peptides (403-423). (D) Peptide pulldowns were performed as described in Figure 3E, with increasing concentrations of NaCl for washes. (E) Peptide pulldowns were performed as described in Figure 3E, except PRC2 was pre-incubated with 37.5 µM of either H3K27M 1-42 or 3x-FLAG peptide in the presence of 40 µM SAM before addition of biotinylated-EZHIP peptide and beads. (F) In vitro PRC2 assays using native PRC2 and oligonucleosome substrate as described in Figure 2B in the presence of increasing concentrations of full-length EZHIP or EZHIP peptide (403-423). (G) Immunoblots of eluates from FLAG M2 immunoprecipitation from nuclear extracts of 293T cells overexpressing FLAG-tagged full length EZHIP or EZHIP fragments (1, 2, 4, 5, 8, 9, 10, 11) as shown in Figure 3H.

**Figure S4:**
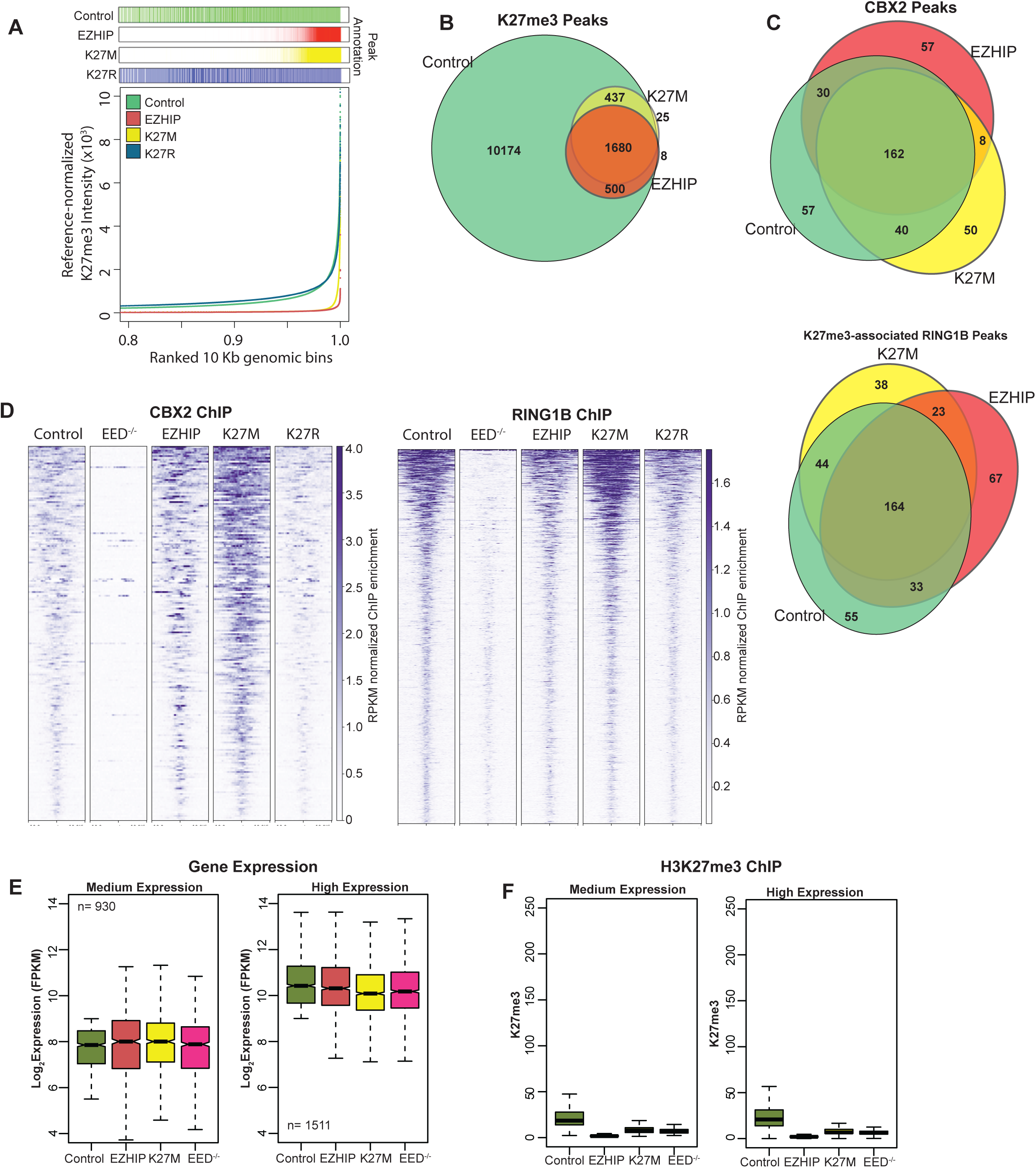
“Residual” peaks of H3K27me3 are sufficient to recruit PRC1 complex. (A) Distribution of reference-normalized H3K27me3 read density across the mouse genome. The genome was binned into 10 kb bins and ranked by their H3K27me3 densities (x-axis). Bins associated with H3K27me3 peaks are shown above the plot. (B) Venn diagram displaying the overlap between H3K27me3 peaks in MEFs overexpressing H3K27M, EZHIP and control cells. (C) Venn diagram displaying the overlap between Cbx2 and Ring1b peaks. (D) Heatmap displaying the RPKM-normalized enrichment of Cbx2 and Ring1b at their corresponding peaks in control cells. (E) Boxplot displaying the expression of medium-(left) and high-expression (right) genes. (F) Boxplot displaying the Rx-normalized H3K27me3 ChIP enrichment at the promoters of genes in E.

**Figure S5:**
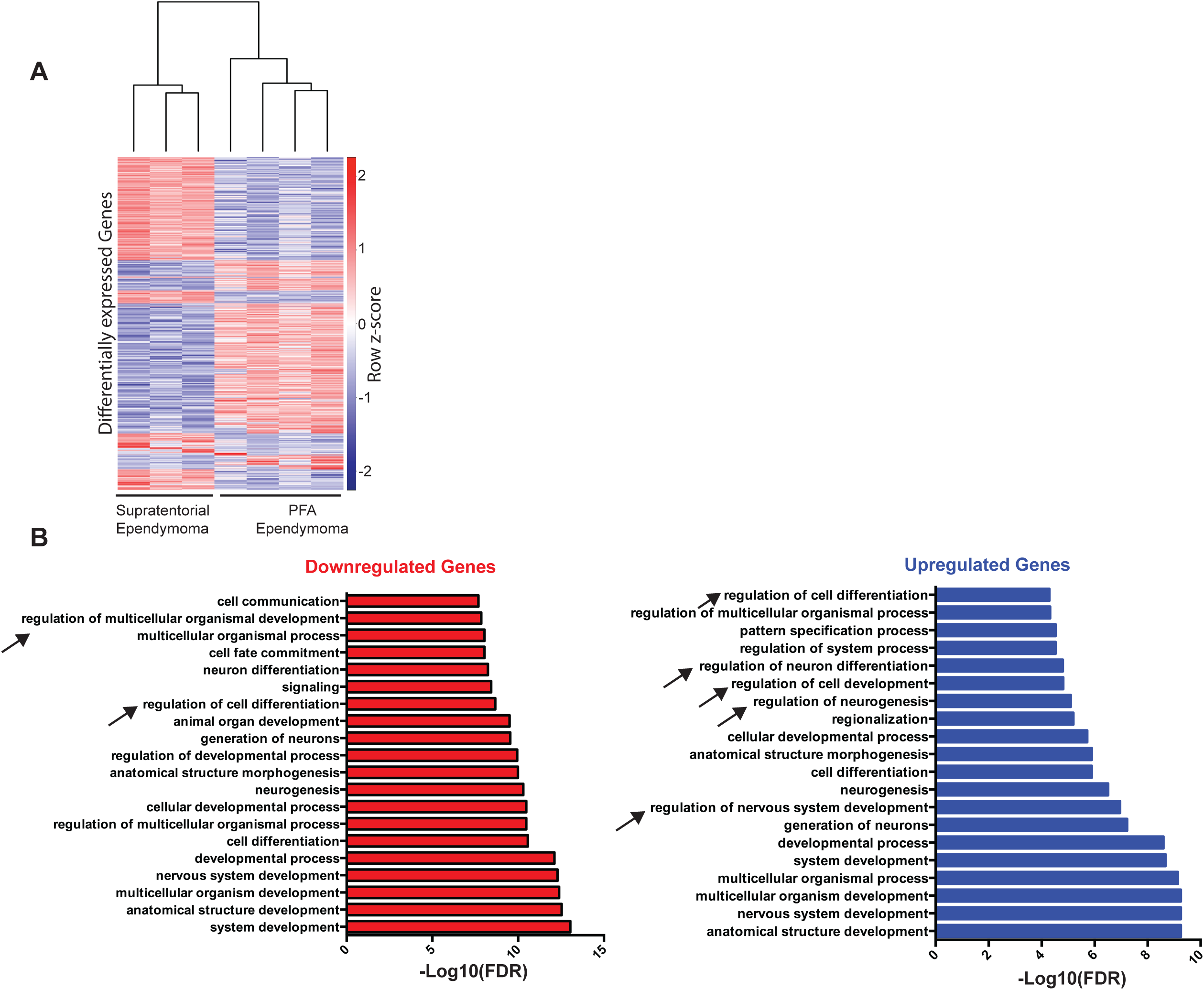
(A) Heatmap displaying the expression profile of differentially expressed genes (>2x difference) in PFA and supratentorial ependymomas. (B) Top 20 GO terms enriched in differentially expressed genes between PFA and supratentorial ependymomas.

## Materials and Methods

### Clonin

Human and mouse EZHIP gene was cloned from genomic DNA isolated from 293T and MEF cells respectively. The protein sequences used in this study are described below.

### Transgenic Cell line generatio

Mouse embryonic fibroablasts (MEFs) used in this study were isolated from E13.5 day embryo that contain loxP sites flanking exons 3-6 of the EED gene (Xie et al., 2014). Human 293T, 293F or Mouse Embryonic Fibroblasts (MEFs) cells were transduced with recombinant lentiviruses produced using pCDH-EF1a-MCS-Puro expression vector. Transduced cells were selected with 1 µg/ml of puromycin for 4 days and harvested for immunoblot analysis after 6-10 days.

### Histone extractio

Cells were lysed in hypotonic lysis buffer (10mM HEPES, 10mM KCl, 1.5mM MgCl2, 0.5mM PMSF, 0.1% TritonX-100. Chromatin pellet was incubated in 0.4N H2SO4 overnight at room temperature. Histones were precipitated from the supernatant using 33% TCA, washed in acetone and resuspended in water.

### Histone derivatization and PTM analysis by nano-LC-M

Acid-extracted histones (5-10 μg) were resuspended in 100 mM ammonium bicarbonate (pH 8), derivatized using propionic anhydride and digested with trypsin as previously described (*26*). The resulting histone peptides were desalted using C18 Stage Tips, dried using a centrifugal evaporator, and reconstituted using 0.1 M acetic acid in preparation for LC-MS analysis. The samples were loaded onto a trap column (75 µm x 2 cm, 3 µm, ReproSil-Pur 120 C18-AQ, Dr. Maisch GmbH) using an Eksigent AS2 autosampler. Histone peptides were resolved on an analytical column (75 µm x 15 cm, 3 µm ReproSil-Pur 120 C18-AQ) using a two-step gradient from 2% ACN to 30% ACN in 0.1% formic acid over 40 min, then from 30% ACN to 95% ACN in 0.1% formic acid over 20 min at a flow rate of 250 nL/min using an Eksigent nanoLC-Ultra 2D. The HPLC was coupled to an Orbitrap Elite mass spectrometer. Full-MS spectra (290-1600 m/z) were collected using a resolution of 60,000 and an AGC target value of 1×10^6^ in the orbitrap. Lock mass calibration in full MS scans was enabled using polysiloxane ions at 371.1012 and 445.1200 m/z. MS/MS data was collected using targeted scans against H3 and H4 peptides in addition to data-dependent scans using collision activated dissociation (CAD) and an AGC target value of 1×10^4^ in the ion trap. Dynamic exclusion was enabled with an exclusion time of 30 s. Singly-charged ions and ions without a detectable charge state were excluded from fragmentation. Mass-to-charge ratios were calculated for each modified form of H3 and H4 peptides, including the H3.3 (27-40) peptide containing the K36M mutation. Data from all detectable charge states were summed. The area under the curve was calculated from extracted ion chromatograms using Xcalibur Qual Browser for each histone peptide mass-to-charge ratio using a 10 ppm mass accuracy cutoff. The area for each modification state of a peptide was normalized against the total peptide signal to give the relative abundance of the histone modification. Each histone sample was injected three times on the MS, and the relative abundance of each peptide modification was averaged across the runs.

### Peptide pulldown assay

25 µL of high capacity Streptavidin-agarose beads containing saturating amounts of biotinylated peptide or no peptide were incubated for 2 hrs with 3 μg recombinant PRC2 purified from SF9 cells overexpressing EZH2, Suz12, EED, and RbAp48. The sequences for each peptide are as follows: H3.3K27M (18-37): KQLATKAARMSAPSTGGVKKYK-biotin; H2B (1-21): PEPSKSAPAPKKGSKKAITKAYK-biotin; EZHIP (403-422): AVRMRASSPSPPGRFFLPIPK-biotin. The incubation was performed with or without 40 μ? SAM in 1mL of 20 mM Tris-HCl pH8, 75mM NaCl, 0.01%NP-40, 0.4 mM PMSF, 1mM β-mercaptoethanol, and 20 μ? nonbiotinylated H3K27me3 stimulator peptide. Following incubation, the beads were washed 4 times in 1mL of the same buffer, except without the stimulator peptide. Bound protein was eluted with acidic glycine (100 mM glycine pH2.5, 150 mM NaCl).

Peptide pulldowns using increasing amounts of NaCl were performed as above with 40 μM SAM present in all samples. Binding was performed at 75 mM NaCl, while washes were done at 75, 150, 250, 500 mM NaCl.

For peptide pulldowns pre-incubated with H3K27M peptide, 3 μg of recombinant PRC2 was pre-incubated for 20 min with 37.5 μmol of nonbiotinylated H3K27M (1-37: ARTKQTARKSTGGKAPRKQLATKAARKSAPATGGVKK) or 37.5 μmol of 3xFLAG peptide. Following pre-incubation, 0.375 μmol of biotinylated-EZHIP peptide was added and incubated for 40 min. Then, 25μL of Streptavidin-agarose beads were added and incubated for an additional 1.5 hrs. All incubations and washing conditions were done as above, using 75mM NaCl.

### Purification of native/recombinant proteins and complexes

Native PRC2 and EZHIP-PRC2 complexes was purified from HeLa and 293T cells expressing FLAG-tagged Ezh2 or FLAG-tagged EZHIP respectively. Nuclei were isolated by resuspending cells in buffer-A (15 mM HEPES pH7.9, 4 mM MgCl2, 10 mM KCl, 1 mM EDTA, 4 mM PMSF). Nuclei were resuspended in buffer-AC (15 mM HEPES pH7.9, 110 mM KCl, 4 mM MgCl_2_, 1 mM EDTA, 0.4 mM PMSF, 1X Protease Inhibitor Cocktail). Nuclear extract was prepared by adding 1/10^th^ volume of saturated ammonium sulfate and ultracentrifugation at 28k RPM for 90 min. Supernatant was dialyzed against FLAG-IP buffer (20mM HEPES pH 7.9, 250 mM KCl, 1mM EDTA, 2 mM β-mercaptoethanol, 0.4 mM PMSF, 0.1% TritonX-100). Nuclear extract was incubated with M2 anti-FLAG affinity gel (Sigma A2220) for 2 hours. Beads were subsequently washed three times with wash buffer (20mM HEPES pH 7.9, 500 mM KCl, 1mM EDTA, 2 mM β-mercaptoethanol, 0.4 mM PMSF, 0.1% TritonX-100). Captured proteins were eluted in 20 mM HEPES pH 7.9, 200 mM KCl, 1 mM EDTA, 10% glycerol, 0.4 mM PMSF, 500 ug/ml 3X-FLAG peptide. Complexes were purified by M2 affinity chromatography from nuclear extract, followed by Mono-Q anion exchange chromatography.

Recombinant PRC2 complex was purified from SF9 cells co-infected with baculoviruses containing human FLAG-tagged Ezh2, Suz12, EED, Aebp2 and RpAp48. Cells were lysed in lysis buffer (15 mM Tris pH8.0, 5 mM MgCl2, 500 mM KCl, 0.5% Triton X-100, 1 mM EDTA, 0.4 mM PMSF, 1X Protease Inhibitor Cocktail, β-mercaptoethanol), followed by M2 affinity purification and anion-exchange chromatography. Similarly, recombinant EZHIP was purified from SF9 cells by M2 affinity purification.

Recombinant EZHIP was purified from *E. coli*-expressing 6x-His-tagged EZHIP overnight at 18°C. Cells were lysed using sonication (20 peak power, 20 sec ON-OFF) for 6 min on ice. EZHIP was purified using Ni-NTA agarose beads, followed by mono-S cation exchange chromatography.

Native oligonucleosomes were purified from Ezh2^-/-^ Mouse Embryonic Fibroblasts. Nuclei were prepared by resuspending 100 million cells in buffer-A and centrifugation at 3000 RPM for 5 min. Nuclei were resuspended in buffer-AP (15mM HEPES pH8, 15mM NaCl, 60mM KCl, 5% Sucrose, 0.5mM Sperimine, 0.15mM Spermidine, 0.4mM PMSF, 1mM β-mercaptoethanol) and treated with 0.2units/ul MNase for 20 min at 37 °C. After quenching with 5 mM EDTA, nuclei were centrifuged at 3000 RPM for 5 min. Nuclei were lysed by resuspension in 10 mM EDTA and 500 mM NaCl. Oligonucleosomes were purified over sucrose gradient (5-30% sucrose, 15mM HEPES pH7.9, 1 mM EDTA, 500 mM NaCl, 5 mM PMSF). Oligonucleosomes in fractions 15-21 ml were concentrated and dialyzed against 15 mM Tris pH 8.0, 100 mM NaCl, 1 mM EDTA, 4 mM PMSF, 10% glycerol.

### Mass spectrometry for protein identification and quantificatio

The eluates from FLAG-Immunoprecipitation were reduced with 10 mM DTT for 30 mins at 56C, alkylated with 50 mM iodoacetamide for 40 mins in the dark at room temperature, and then diluted into 4 volumes of 50 mM Tris pH 8 with 2 mM CaCl2 for overnight digestion with trypsin (1 ug) at 37C. After desalting with C18 stage tips, the peptides were analyzed by LC-MS/MS with an EasyLC1000 nanoLC system (Thermo) and a Fusion Orbitrap mass spectrometer (Thermo). MS was performed over a 95 min LC gradient with a 2 second cycle of one full MS scan in the orbitrap followed by DDA MS/MS scans in the ion trap on the most abundant precursor ions fragmented by HCD at 27NCE. Dynamic exclusion was set to 40 sec.

Mass spec raw files were processed by ProteomeDiscoverer software (Thermo). Protein abundance ratios were expressed as specific FLAG pulldown divided by mock pulldown. PD results were exported as text files to Excel. Proteins with at least 2 unique peptides and log2 ratios (FLAG/mock) greater than 4 were considered as hits.

### Histone Methyltransferase assay

For a typical PRC2 reaction, 200 nM oligonucleosome or 50 µM H3 (18-37) peptide substrate was incubated with 20 nM PRC2 complex, 4 µM S-adenosyl Methionine (1 µM ^3^H-SAM (Perkin Elmer); 3 µM cold SAM) and 20 µM H3K27me3 peptide in 2X reaction buffer (50 mM Tris pH 8.0, 4 mM MgSO4, 10 mM DTT and 8 mM PMSF) for 60 min. For scintillation couting, reaction was spotted on a phosphocellulose filter (Whatman p81) and dried for 10 min. Filters were washed 3x with 0.1 M NaHCO_3_ for 5 min, rinsed with acetone and dried for 10 min. Scinitillation couting was performed using Tri-Carb 2910 TR liquid scintillation analyzer (Perkin Elmer). Counts were corrected for background using reaction without substrate. For flurography analysis, reaction mixture was resolved on SDS-PAGE gel, stained with coommassie blue stain and incubated in Amersham Amplify Flurography Reagent (GE Healthcare) and dried under vacuum. Films capturing flurographic signal were developed after 24 hours of exposure.

### Chromatin Immunoprecipitation

Cells were cross-linked using 1% paraformaldehyde for 5min at room-temperature and quenched with 200mM glycine. ∼20 million cells were lysed by resuspending in digestion buffer (20 mM Tris pH 7.6, 1mM CaCl2, 0.25% TritonX-100, 5mM PMSF, 1x Protease inhibitor cocktail). Chromatin was treated with 10 units MNase/ million cells for 10 min at 37°C and quenched by adding 5mM EDTA. Chromatin was solubilized by sonication using covaris (120 Peak incidental power, 5% duty factor, 200 cycles/burst) for 3 minutes. Chromatin was dialyzed against RIPA buffer (10mM Tris pH8, 0.1% SDS, 0.1% Na-DOC, 1 mM EDTA, 1% TritonX-100) for 2 hours. Spike-in (293T or S2 for MEFs) chromatin was added at 1:40 dilution and incubated with antibody overnight. Chromatin bound to antibody was captured by dynabeads for 3 hours, washed (3x RIPA, 3x RIPA + 300mM NaCl, 2x RIPA-LiCl for 5 min each) and eluted in 10mM Tris, 1 mM EDTA, 1% SDS. Eluted DNA was reverse crosslinked overnight at 65oC, treated with proteinase-K and purified using PCR purification kit. Illumina sequencing libraries were generated using NEB Ultra kit.

### ChIP-Seq analysi

ChIP-Seq data was analyzed as previously described (Lu et al. 2016). Briefly, reads that passed quality score were aligned to mouse mm9 and reference genomes (dm6 or hg19) using bowtie1 with default parameters. Sample normalization factor was determined as Rx factor = 10^6^/(reads mapped to reference genome) or RPKM factor = 10^6^/total aligned reads. Peaks were determined using Mosaics-HMM (broad peaks) for H3K27me3, CBX2 and Ring1b ChIPs (FDR= 0.1, maxgap= 10,000, minsize= 1000. Thres= 30, 8 and 20 respectively). Bigwig files were generated using deeptools and visualized using IGV genome browser. Published ChIP-Seq data from Stafford et al. was analyzed similarly. Statistical analysis and graphs were generated using R.

Published ChIP-Seq data from Bayliss et al. was analyzed as previously described. However, peaks were called using mosaicsHMM for all samples with the following parameters: FDR= 0.1, maxgap= 20,000, minsize= 400. Thres= 15 for supratentorial samples, whereas thres= 20 for posterior-fossa samples to adjust for broad, low-intensity verses sharp high-intensity peaks respectively.

### RNA Preparation

RNA was purified from MEFs using Trizol Reagent. RNA purified from FFT cells was spiked into each sample at a 1:50 ratio. The RNA was DNase treated using Turbo DNase Treatment (Ambion AM1907). Then, the RNA was rRNA-depleted using NEBNext rRNA Depletion Kit E6310L. Libraries were generated using NEB E7530.

### RNA-Seq analysi

Reads that passed quality filter were aligned and expression values were determined using RSEM. Differentially expressed genes were determined using EBSeq with a fold-change cutoff of 2-fold and posterior probability of differential expression >= 0.95. Graphs and statistical analyses were performed using R.

### Antibodies

H3K9me3: Active Motif 39161

H3K27ac: Active Motif 39133

H3K27me1: Millipore 07-448

H3K27me2: Cell Signaling d18C8

H3K27me3: Cell Signaling C36B11

H3 general: Proteintech S2900-1

H4 general: Proteintech S2901-2

H3K36me3: Active motif 61101

H3K36me2: Cell Signaling 2901S

FLAG: M2 Sigma Aldrich F1804

EZHIP: atlas hpa004003

EZH2: BD Biosciences 612666

EED: Active Motif clone 41D 61203 SUZ12: Cell Signaling D39F6 RBBP4: Proteintech 20364-I-ap RBBP4/6: LP bio AR-01-0178-200

Ring1b: Active motif 39663

Jarid2: Cell signaling 13594S

Phf19: Cell signaling 77271S

Brd4: bethyl a301-985a50

Cbx2: bethyl a302-524a

### Sequences

#### Human EZHIP

MDYKDDDDKGAGMATQSDMEKEQKHQQDEGQGGLNNETALASGDACGTGNQDPAASVTTVSSQASPSGGAALSSSTAGSSAAAATSAAIFITDEASGLPIIAAVLTERHSDRQDCRSPHEVFGCVVPEGGSQAAVGPQKATGHADEHLAQTKSPGNSRRRKQPCRNQAAPAQKPPGRRLFPEPLPPSSPGFRPSSYPCSGASTSSQATQPGPALLSHASEARPATRSRITLVASALRRRASGPGPVIRGCTAQPGPAFPHRATHLDPARLSPESAPGPARRGRASVPGPARRGCDSAPGPARRGRDSAPVSAPRGRDSAPGSARRGRDSAPGPALRVRTARSDAGHRSTSTTPGTGLRSRSTQQRSALLSRSLSGSADENPSCGTGSERLAFQSRSGSPDPEVPSRASPPVWHAVRMRASSPSPPGRFFLPIPQQWDESSSSSYASNSSSPSRSPGLSPSSPSPEFLGLRSISTPSPESLRYALMPEFYALSPVPPEEQAEIESTAHPATPPEP

#### Mouse EZHIP

MASSSSPERGLEALRDTDESEGEAPGPSGPRGRGGPSGAGSALRLRSLEAEMAAACVTSTAGEDLGTFSEPGSQHGDPEGGGGPDLELGHARPMMRSQRELGLTPKGGGKADQGGKGRKGGSGSPPHTKSSRKREQPNPNRSLMAQGAAGPPLPGARGSPAMPQPESSLSPRPDQSHHFDFPVGNLEAPGPTLRSSTSQGSGSTPVPEALRCAESSRAESDQSSPAGRELRQQASPRAPDDDDDGDGGPDPRGSGTPEGWVLRSGVVPFGRRSSASEVSPEEVRPEAQCTGWNLRPRPRSSASAVSPEARPKAQSAGRNLRPRPRSSASVVSPEARPKAQSAGRNLRPRPRSSASVVSPEARPEAQSAGRNLRPRATPRVPVAPSSTTRSSSDRGSSRAPRSRSRSRSCSTPRLGSDHQRSRKIKMRLDLQVDREPESEAEQEEQELESEPGPSSRPQASRSSSRFAVPGRSSLAAEDSPPRRPVRMRASSPSPPGRLYPLPKHYFEGVHSPSSSSSESSSVSSSHSPLNKAPDPGSSPPLSSLSGPNPFWLALIADLDNLDSSSPRVPGEEIEAAPHTREEEDKKCRG

#### EZHIP-H3.3 fusion protein

MARTKQTARKSTGGKAPRKQLATKAVRMRASSPSPPGRFFLPRPGTVALREIRRYQKSTELLIRKLPFQRLVREIAQDFKTDLRFQSAAIGALQEASEAYLVGLFEDTNLCAIHAKRVTIMPKDIQLARRIRGERAAAAGGDYKDDDDKSAAGGYPYDVPDYA

